# Oligomannosylation and MAN1A1 expression associate strongly with a subset of human cancer types

**DOI:** 10.1101/2021.05.08.443254

**Authors:** Sayantani Chatterjee, Rebeca Kawahara, Julian Ugonotti, Ling Y. Lee, Arun Everest-Dass, Morten Thaysen-Andersen

## Abstract

Aberrant protein glycosylation is a prominent cancer feature. While many tumour-associated glycoepitopes have been reported, advances in glycoanalytics continue to uncover new associations between glycoproteins and cancer. Guided by a comprehensive literature survey suggesting that oligomannosylation (Man_5-9_GlcNAc_2_, M5-M9) is a widespread albeit poorly studied glyco-signature in human cancers, we here re-visit a valuable compilation of nearly 500 LC-MS/MS *N*-glycomics datasets acquired across 11 human cancer types to systematically test for oligomannose-cancer associations. Firstly, our quantitative glycomics data obtained across 34 cancerous cell lines demonstrated that oligomannosylation, particularly the under-processed M7-M9, is a strong pan-cancer feature. We then showed cell surface expression of oligomannosidic epitopes in the promyelocytic leukemic HL-60 cell line using concanavalin A-based flow cytometry. In keeping with literature, our quantitative glycomics data of tumour and matching control tissues and new MALDI-MS imaging data of tissue microarrays showed a strong cancer-associated elevation of oligomannosylation in both basal cell (*p* = 1.78 x 10^-12^) and squamous cell (*p* = 1.23 x 10^-11^) skin cancer and colorectal cancer (*p* = 8.0 x 10^-4^). The glycomics data also indicated that few cancer types including gastric and liver cancer exhibit unchanged or reduced oligomannose levels, observations also supported by literature and MALDI-MSI. Finally, data from cancer repositories indicated that three α1,2-mannosidases dictate oligomannose expression in cancer cells, and further suggested that deleterious mutations and reduced expression of MAN1A1 are key contributors to the cancer-associated oligomannose elevation. Collectively, these findings open hitherto unexplored avenues for the development of new cancer biomarkers and therapeutic targets.

## 1. Introduction

Protein glycosylation, the addition of complex carbohydrates (glycans) to polypeptides, is a ubiquitous and energy-demanding post-translational modification important for a plethora of inter- and intracellular processes [1–5].

Asparagine (Asn, *N*)-linked glycans display a remarkable molecular heterogeneity despite being assembled from only few different monosaccharide building blocks including mannose (Man), galactose (Gal), fucose (Fuc), glucose (Glc), *N*-acetylglucosamine (GlcNAc) and *N*-acetylneuraminic acid (NeuAc) [5]. The template-less *N*-glycan biosynthesis begins in the endoplasmic reticulum (ER) by the transfer of immature Glc-capped glycan precursors to Asn residues located in consensus sequences of acceptor polypeptides and continues in the Golgi compartments. The biosynthesis involves successive glycan processing steps catalysed by dozens of glycosyltransferases and glycoside hydrolases [6–8]. The resulting *N*-glycans that inherently display an extensive molecular heterogeneity are commonly classified as oligomannosidic-, hybrid- or complex-type *N*-glycans [9–12], **Figure 1A**. Recently, truncated *N*-glycans not fitting into these three classes were reported from various human biospecimens including the inflammation- and cancer-associated paucimannosidic-(Man_1-3_GlcNAc_2_Fuc_0-1_) and the even shorter chitobiose core-(GlcNAc_1-2_Fuc_0-1_) type *N*-glycans [12–16]. Several glycan species fit into each glycan type; for example, the M5-M9 glycans (Man_5-9_GlcNAc_2_), the focus of this study, belong to the oligomannosidic-type *N*-glycans.

**Figure 1.**
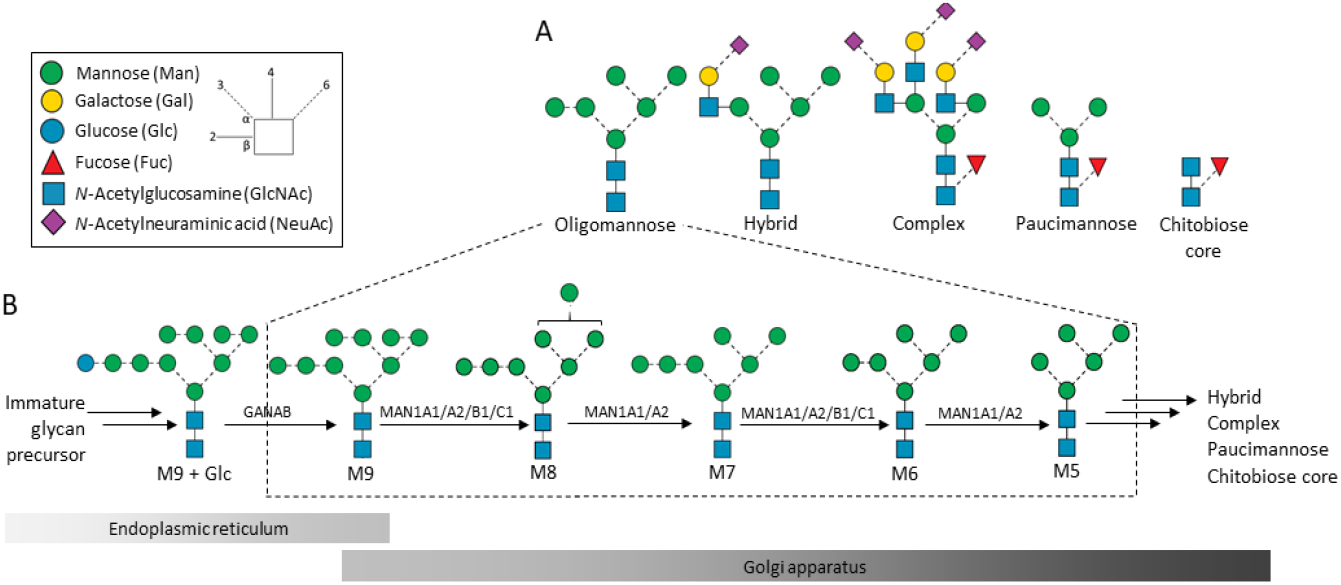
Structures, nomenclature and biosynthesis of protein oligomannosylation. (**A**) Oligomannose, one of five *N*-glycan types in human glycobiology, remains understudied in cancer research despite forming a large and vital part of the *N*-glycome. (**B**) Oligomannosidic *N*-glycans are formed via enzymatic processing early in the biosynthetic pathway from immature Glc-capped glycan precursors in the endoplasmic reticulum (ER). The GANAB-mediated trimming of M9+Glc to M9 is followed by the processing by multiple ER- and Golgi-resident α1,2-mannosidases (MAN1A1, MAN1A2, MAN1B1, and MAN1C1) that successively trim M9 to M5. Each of the oligomannosidic glycans is known to exist as multiple isomers; only commonly observed isomers are depicted in this figure, see **Supporting Data 1** for more. The glycan processing may in cancer and other cell types terminate during the oligomannose trimming or continue to form more processed *N*-glycan types generating an extensive micro-heterogeneity typically observed for glycoproteins. Insert: key to monosaccharide symbols and linkages [1].

Oligomannosidic *N*-glycans are synthesised early in the biosynthetic pathway, **Figure 1B**. Several ER- and Golgi-resident α1,2-mannosidases catalyse the successive M9-to-M5 glycan processing. Specifically, mannosyl-oligosaccharide 1,2-α-mannosidase 1A (MAN1A1, UniProtKB, P33908), mannosyl-oligosaccharide 1,2-α-mannosidase IB (MAN1A2, O60476), endoplasmic reticulum mannosyl-oligosaccharide 1,2-α-mannosidase (MAN1B1, Q9UKM7) and mannosyl-oligosaccharide 1,2-α-mannosidase 1C (MAN1C1, Q9NR34) are known to trim the outer α1,2-Man residues of nascent glycoproteins [17, 18]. While these first processing steps play recognised roles in glycoprotein folding, maturation and trafficking, oligomannosylation is also involved in a host of less studied extracellular functions including cell-cell and cell-extracellular matrix communication relevant for cancer processes [2, 19–22].

Decades of intense research efforts have provided robust evidence supporting that altered protein glycosylation is an inherent feature of malignant transformation and other cancer traits, and have unearthed many glycan structures and glycoepitopes that play important roles in cancer [23]. Excellent reviews have summarised our current knowledge of the aberrant protein glycosylation underpinning cancer [23–28]. Altered expression patterns of fucosylation e.g. core and Lewis^X^ [29, 30], sialylation e.g. α2,3- and α2,6-NeuAc [31, 32] and glycan antennary branching e.g. bisecting β1,4-GlcNAc [33, 34] are prominent examples of glycan features repeatedly associated with cancer.

Despite forming a considerable part of the *N*-glycome, the mannose-terminating *N*-glycans comprising both the oligomannosidic- and paucimannosidic-type have received comparably less attention in cancer research [22]. We recently used quantitative glycomics to systematically document that paucimannosylation is an overlooked feature in many cancer types including in brain (glioblastoma and neuroblastoma), blood (acute lymphocytic leukaemia, ALL; acute monocytic leukaemia, AML; acute promyelocytic leukaemia, APL; chronic lymphocytic leukemia, CLL), bladder (BlaCa), melanoma and non-melanoma (basal cell carcinoma, BCC and squamous cell carcinoma, SCC) skin, breast (BC), hepatocellular carcinoma (HCC, liver), lung, ovarian (OvC), gastric (GC), colorectal (CRC) and prostate cancer (PCa) [35]. These findings were supported by quantitative glycomics data from nearly 500 LC-MS/MS datasets acquired over 10 years across laboratories around the world from a variety of cancer specimens including cultured human cancer cells and tissues from cancer patients and matching controls using a uniform analytical platform. The paucimannosidic-centric study, however, did not explore the wealth of other valuable information found in this compilation of content-rich *N*-glycomics datasets.

Guided by an initial literature survey suggesting that associations exist between oligomannosylation and human cancers, we have herein re-interrogated our unique collection of *N*-glycomics datasets to systematically test for oligomannose-cancer relationships across human cancer types. It transpires from this meta-analysis and supporting data from new matrix-assisted laser desorption/ionization mass spectrometry imaging (MALDI-MSI) and lectin flow cytometry analyses and enzyme expression and mutation data retrieved from well-curated cancer repositories that elevated oligomannosylation and reduced MAN1A1 expression are strong features of a subset of human cancers including skin and colorectal cancer.

## 2. Methods

### 2.1. Literature survey

The glycomics and glycobiology literature was comprehensively surveyed in attempts to document a link between oligomannose and human cancer using the PubMed (https://pubmed.ncbi.nlm.nih.gov/) and Google Scholar (https://scholar.google.com.au/) search engines. Combinations of search keywords were employed including “high-mannose” (with/without hyphen), “extended mannose”, “oligomannose”, “human”, “cancer” and “carcinoma”. Other keywords including “*N*-glycan”, “glycoprofiling”, and “*N*-glycome profiling” were used to retrieve additional literature. All types of human cancers, biological samples, and experimental and analytical methods were considered in the literature search. Only original contemporary research papers published from 2000-2021 were considered and included in **Table 1**. Relevant reviews and associated research papers were also surveyed to ensure that the existing literature was exhaustively captured by our search efforts.

**Table 1.**
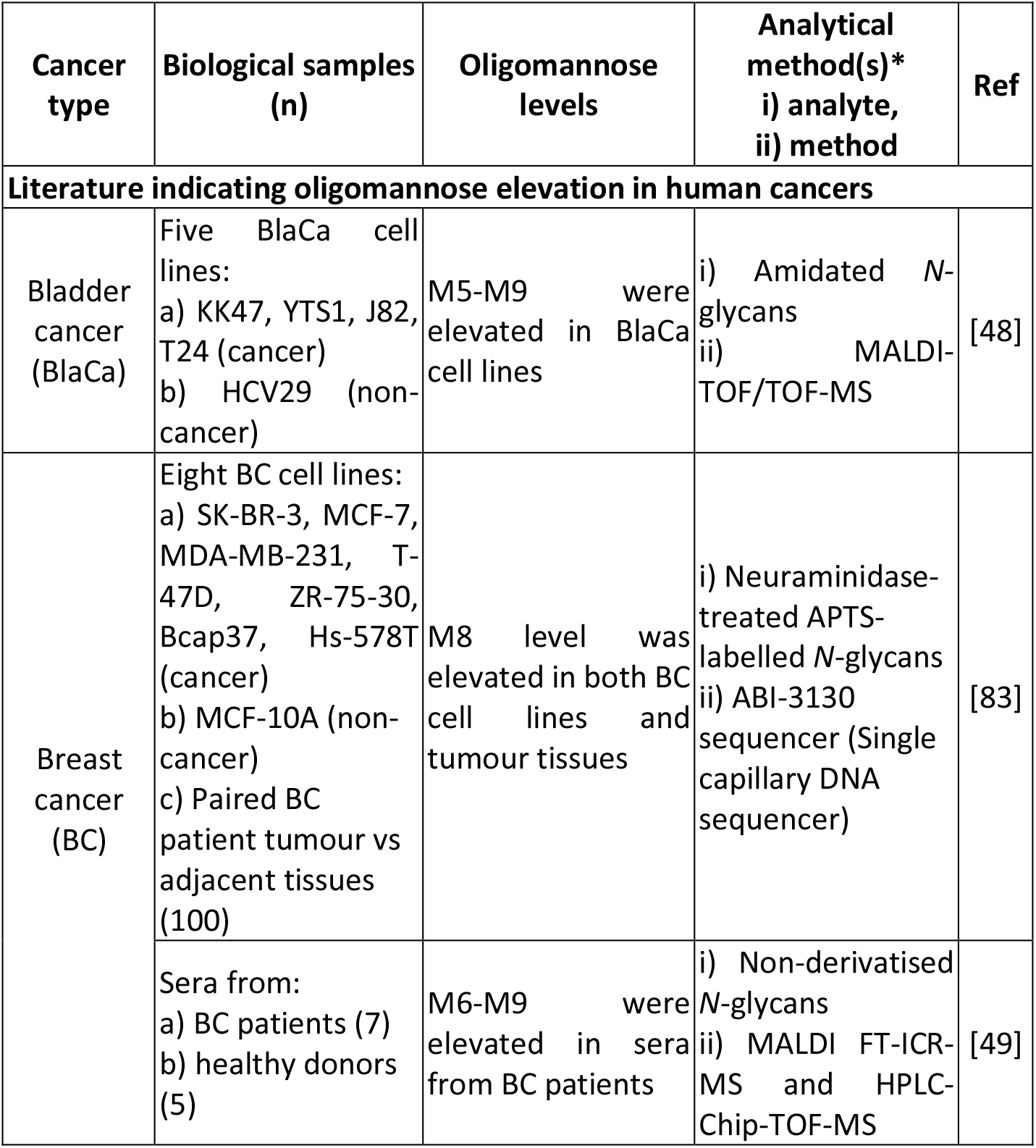
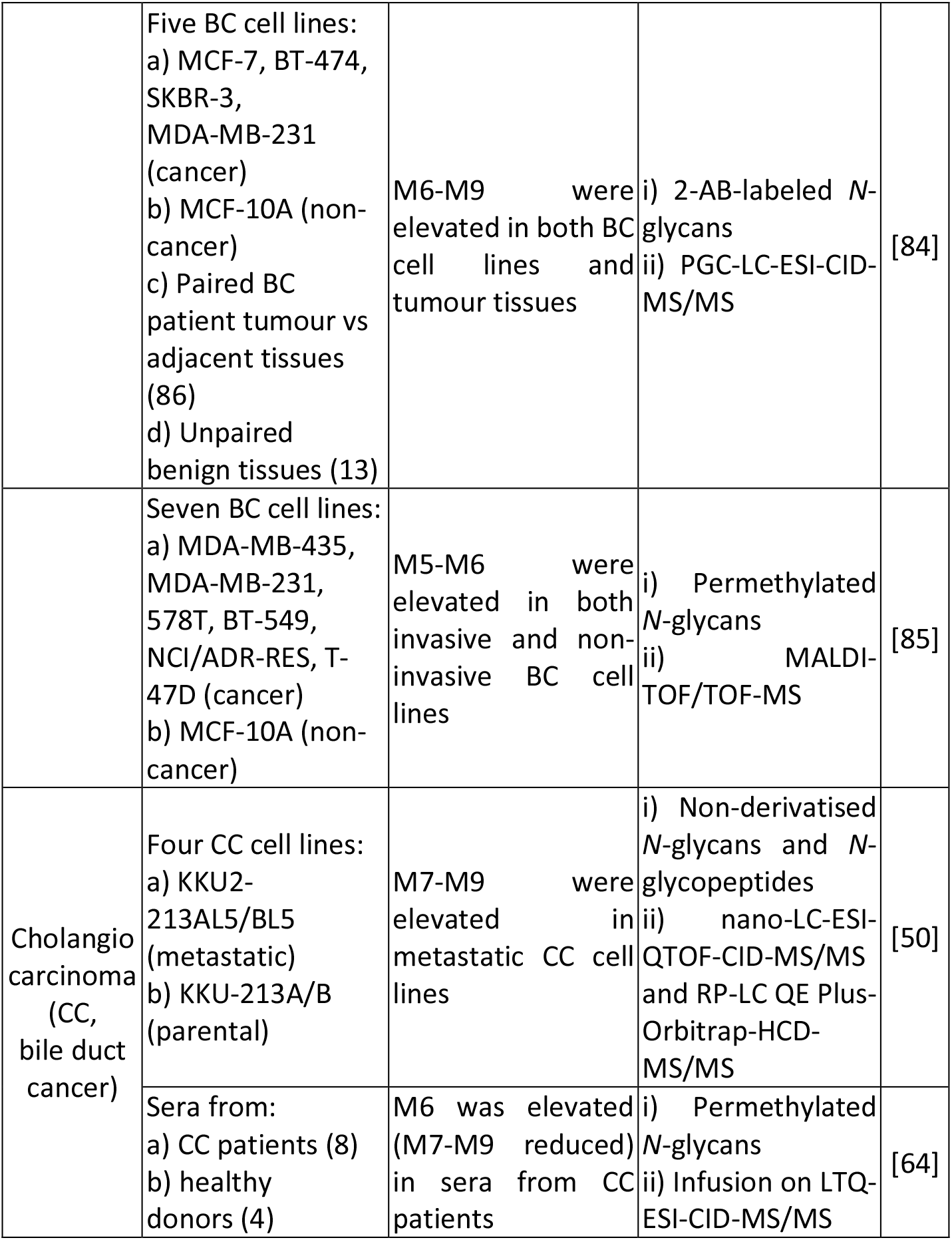
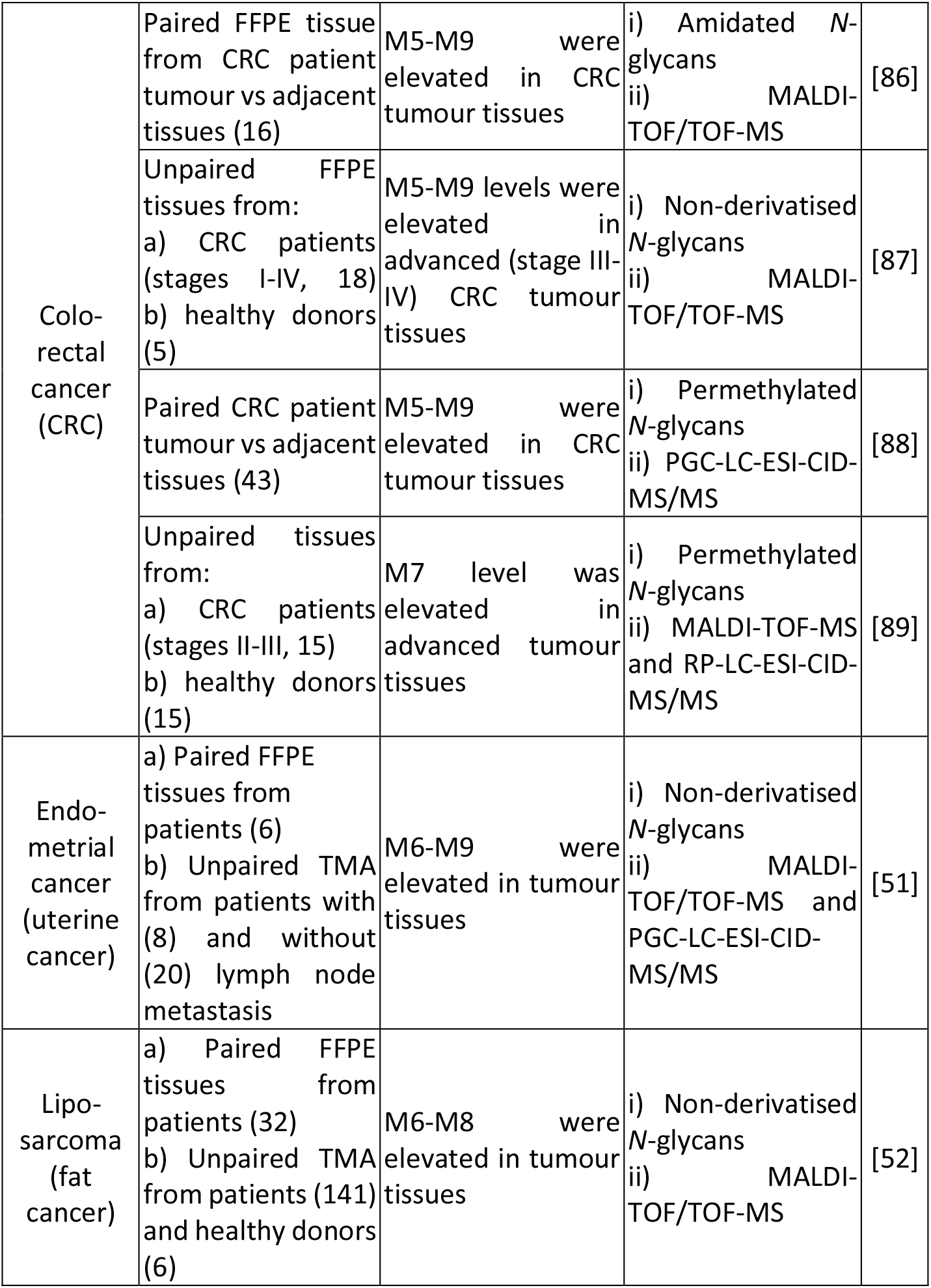
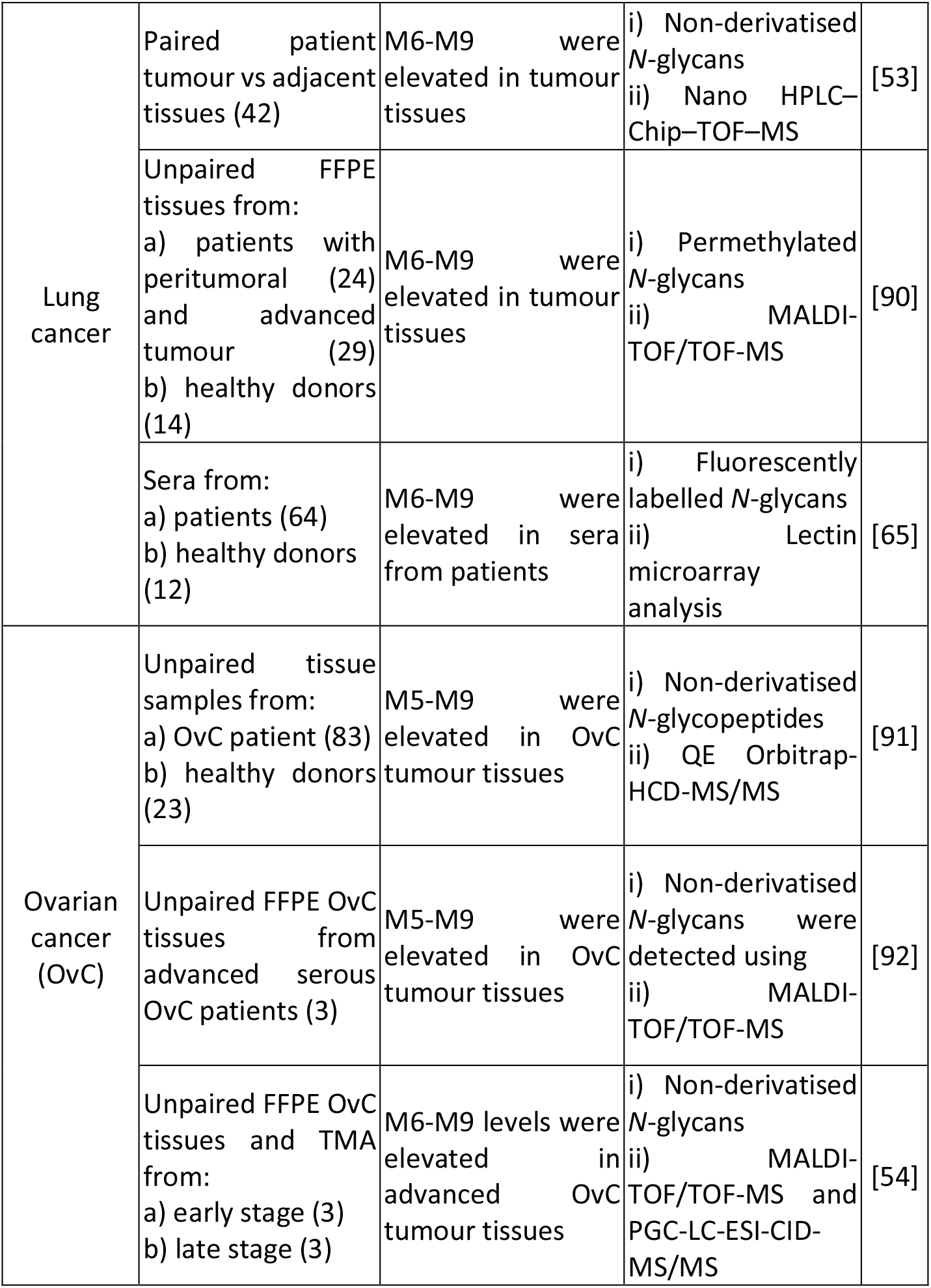
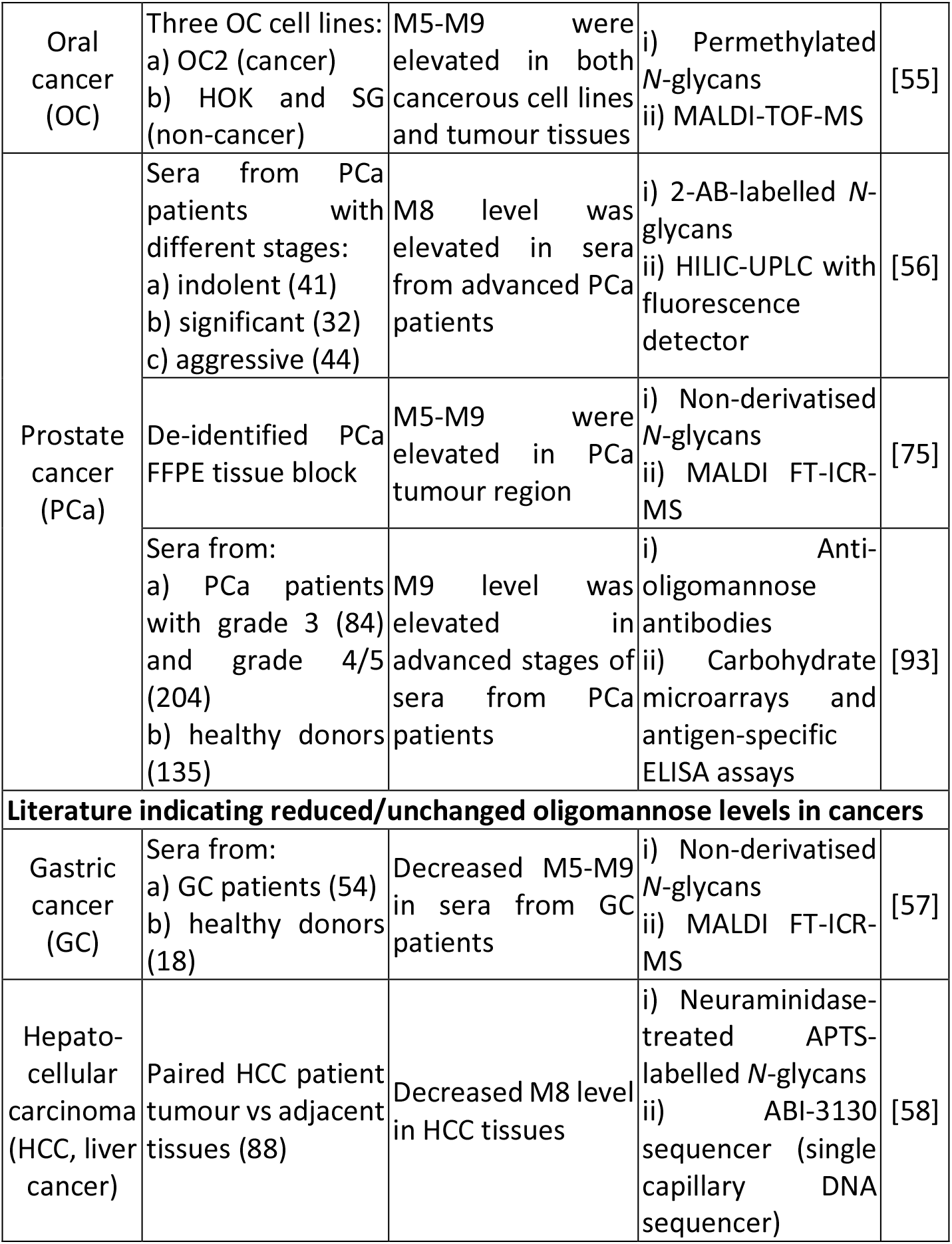
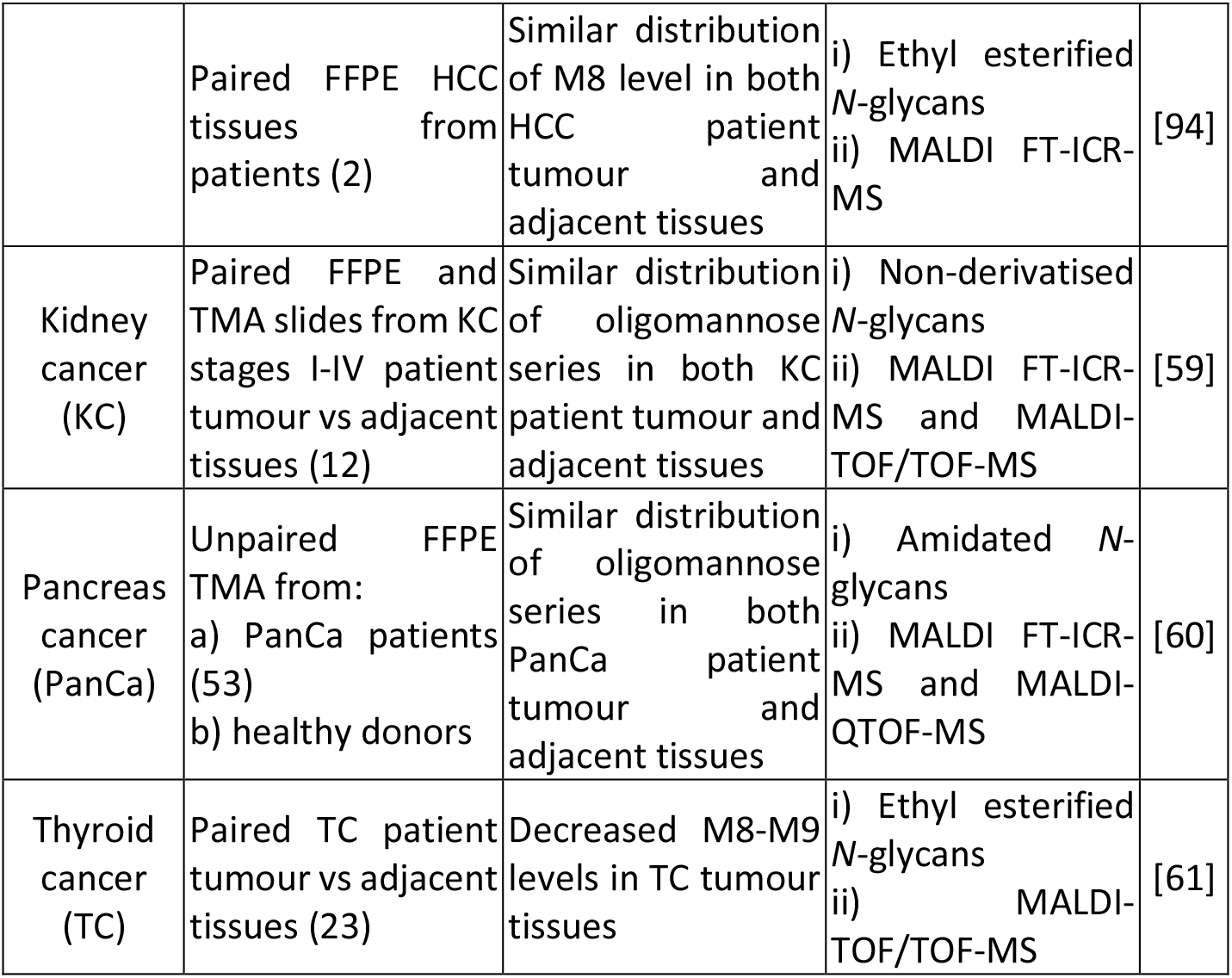
Literature survey exploring a possible link between oligomannose and human cancers. **\***All *N*-glycans were liberated from their carrier proteins using peptide:*N*-glycosidase F. 2-AB: 2-aminobenzoic acid; APTS: 8-amino-1,3,6-pyrenetrisulfonic acid; FFPE: formalin-fixed paraffin-embedded; TMA: tissue microarray.

| Cancer type | Biological samples (n) | Oligomannose levels | Analytical method(s)*<br>i) analyte,<br>ii) method | Ref |
| --- | --- | --- | --- | --- |
| **Literature indicating oligomannose elevation in human cancers** |  |  |  |  |
| Bladder cancer (BlaCa) | Five BlaCa cell lines:<br>a) KK47, YTS1, J82, T24 (cancer)<br>b) HCV29 (non-cancer) | M5-M9 were elevated in BlaCa cell lines | i) Amidated *N*-glycans<br>ii) MALDI-TOF/TOF-MS | [48] |
| Breast cancer (BC) | Eight BC cell lines:<br>a) SK-BR-3, MCF-7, MDA-MB-231, T-47D, ZR-75-30, Bcap37, Hs-578T (cancer)<br>b) MCF-10A (non-cancer)<br>c) Paired BC patient tumour vs adjacent tissues (100) | M8 level was elevated in both BC cell lines and tumour tissues | i) Neuraminidase-treated APTS-labelled *N*-glycans<br>ii) ABI-3130 sequencer (Single capillary DNA sequencer) | [83] |
|  | Sera from:<br>a) BC patients (7)<br>b) healthy donors (5) | M6-M9 were elevated in sera from BC patients | i) Non-derivatised *N*-glycans<br>ii) MALDI FT-ICR-MS and HPLC-Chip-TOF-MS | [49] |
|  | Five BC cell lines:<br>a) MCF-7, BT-474, SKBR-3, MDA-MB-231 (cancer)<br>b) MCF-10A (non-cancer)<br>c) Paired BC patient tumour vs adjacent tissues (86)<br>d) Unpaired benign tissues (13) | M6-M9 were elevated in both BC cell lines and tumour tissues | i) 2-AB-labeled *N*-glycans<br>ii) PGC-LC-ESI-CID-MS/MS | [84] |
|  | Seven BC cell lines:<br>a) MDA-MB-435, MDA-MB-231, 578T, BT-549, NCI/ADR-RES, T-47D (cancer)<br>b) MCF-10A (non-cancer) | M5-M6 were elevated in both invasive and non-invasive BC cell lines | i) Permethylated *N*-glycans<br>ii) MALDI-TOF/TOF-MS | [85] |
| Cholangio carcinoma (CC, bile duct cancer) | Four CC cell lines:<br>a) KKU2-213AL5/BL5 (metastatic)<br>b) KKU-213A/B (parental) | M7-M9 were elevated in metastatic CC cell lines | i) Non-derivatised *N*-glycans and *N*-glycopeptides<br>ii) nano-LC-ESI-QTOF-CID-MS/MS and RP-LC QE Plus-Orbitrap-HCD-MS/MS | [50] |
|  | Sera from:<br>a) CC patients (8)<br>b) healthy donors (4) | M6 was elevated (M7-M9 reduced) in sera from CC patients | i) Permethylated *N*-glycans<br>ii) Infusion on LTQ-ESI-CID-MS/MS | [64] |
| Colo-rectal cancer (CRC) | Paired FFPE tissue from CRC patient tumour vs adjacent tissues (16) | M5-M9 were elevated in CRC tumour tissues | i) Amidated N-glycans<br>ii) MALDI-TOF/TOF-MS | [86] |
|  | Unpaired FFPE tissues from:<br>a) CRC patients (stages I-IV, 18)<br>b) healthy donors (5) | M5-M9 levels were elevated in advanced (stage III-IV) CRC tumour tissues | i) Non-derivatised N-glycans<br>ii) MALDI-TOF/TOF-MS | [87] |
|  | Paired CRC patient tumour vs adjacent tissues (43) | M5-M9 were elevated in CRC tumour tissues | i) Permethylated N-glycans<br>ii) PGC-LC-ESI-CID-MS/MS | [88] |
|  | Unpaired tissues from:<br>a) CRC patients (stages II-III, 15)<br>b) healthy donors (15) | M7 level was elevated in advanced tumour tissues | i) Permethylated N-glycans<br>ii) MALDI-TOF-MS and RP-LC-ESI-CID-MS/MS | [89] |
| Endo-metrial cancer (uterine cancer) | a) Paired FFPE tissues from patients (6)<br>b) Unpaired TMA from patients with (8) and without (20) lymph node metastasis | M6-M9 were elevated in tumour tissues | i) Non-derivatised N-glycans<br>ii) MALDI-TOF/TOF-MS and PGC-LC-ESI-CID-MS/MS | [51] |
| Lipo-sarcoma (fat cancer) | a) Paired FFPE tissues from patients (32)<br>b) Unpaired TMA from patients (141) and healthy donors (6) | M6-M8 were elevated in tumour tissues | i) Non-derivatised N-glycans<br>ii) MALDI-TOF/TOF-MS | [52] |
| Lung cancer | Paired patient tumour vs adjacent tissues (42) | M6-M9 were elevated in tumour tissues | i) Non-derivatised *N*-glycans<br>ii) Nano HPLC–Chip–TOF–MS | [53] |
|  | Unpaired FFPE tissues from:<br>a) patients with peritumoral (24) and advanced tumour (29)<br>b) healthy donors (14) | M6-M9 were elevated in tumour tissues | i) Permethylated *N*-glycans<br>ii) MALDI-TOF/TOF-MS | [90] |
|  | Sera from:<br>a) patients (64)<br>b) healthy donors (12) | M6-M9 were elevated in sera from patients | i) Fluorescently labelled *N*-glycans<br>ii) Lectin microarray analysis | [65] |
| Ovarian cancer (OvC) | Unpaired tissue samples from:<br>a) OvC patient (83)<br>b) healthy donors (23) | M5-M9 were elevated in OvC tumour tissues | i) Non-derivatised *N*-glycopeptides<br>ii) QE Orbitrap-HCD-MS/MS | [91] |
|  | Unpaired FFPE OvC tissues from advanced serous OvC patients (3) | M5-M9 were elevated in OvC tumour tissues | i) Non-derivatised *N*-glycans were detected using<br>ii) MALDI-TOF/TOF-MS | [92] |
|  | Unpaired FFPE OvC tissues and TMA from:<br>a) early stage (3)<br>b) late stage (3) | M6-M9 levels were elevated in advanced OvC tumour tissues | i) Non-derivatised *N*-glycans<br>ii) MALDI-TOF/TOF-MS and PGC-LC-ESI-CID-MS/MS | [54] |
| Oral cancer (OC) | Three OC cell lines:<br>a) OC2 (cancer)<br>b) HOK and SG (non-cancer) | M5-M9 were elevated in both cancerous cell lines and tumour tissues | i) Permethylated *N*-glycans<br>ii) MALDI-TOF-MS | [55] |
| Prostate cancer (PCa) | Sera from PCa patients with different stages:<br>a) indolent (41)<br>b) significant (32)<br>c) aggressive (44) | M8 level was elevated in sera from advanced PCa patients | i) 2-AB-labelled *N*-glycans<br>ii) HILIC-UPLC with fluorescence detector | [56] |
|  | De-identified PCa FFPE tissue block | M5-M9 were elevated in PCa tumour region | i) Non-derivatised *N*-glycans<br>ii) MALDI FT-ICR-MS | [75] |
|  | Sera from:<br>a) PCa patients with grade 3 (84) and grade 4/5 (204)<br>b) healthy donors (135) | M9 level was elevated in advanced stages of sera from PCa patients | i) Anti-oligomannose antibodies<br>ii) Carbohydrate microarrays and antigen-specific ELISA assays | [93] |
| **Literature indicating reduced/unchanged oligomannose levels in cancers** |  |  |  |  |
| Gastric cancer (GC) | Sera from:<br>a) GC patients (54)<br>b) healthy donors (18) | Decreased M5-M9 in sera from GC patients | i) Non-derivatised *N*-glycans<br>ii) MALDI FT-ICR-MS | [57] |
| Hepato-cellular carcinoma (HCC, liver cancer) | Paired HCC patient tumour vs adjacent tissues (88) | Decreased M8 level in HCC tissues | i) Neuraminidase-treated APTS-labelled *N*-glycans<br>ii) ABI-3130 sequencer (single capillary DNA sequencer) | [58] |
|  | Paired FFPE HCC tissues from patients (2) | Similar distribution of M8 level in both HCC patient tumour and adjacent tissues | i) Ethyl esterified *N*-glycans<br>ii) MALDI FT-ICR-MS | [94] |
| Kidney cancer (KC) | Paired FFPE and TMA slides from KC stages I-IV patient tumour vs adjacent tissues (12) | Similar distribution of oligomannose series in both KC patient tumour and adjacent tissues | i) Non-derivatised *N*-glycans<br>ii) MALDI FT-ICR-MS and MALDI-TOF/TOF-MS | [59] |
| Pancreas cancer (PanCa) | Unpaired FFPE TMA from:<br>a) PanCa patients (53)<br>b) healthy donors | Similar distribution of oligomannose series in both PanCa patient tumour and adjacent tissues | i) Amidated *N*-glycans<br>ii) MALDI FT-ICR-MS and MALDI-QTOF-MS | [60] |
| Thyroid cancer (TC) | Paired TC patient tumour vs adjacent tissues (23) | Decreased M8-M9 levels in TC tumour tissues | i) Ethyl esterified *N*-glycans<br>ii) MALDI-TOF/TOF-MS | [61] |

### 2.2. Re-interrogation of cancer *N*-glycomics datasets

The biological samples, sample handling and data acquisition methods used to generate the compilation of LC-MS/MS glycomics datasets reinterrogated with this study have been exhaustively described in Chatterjee et al., 2019 [35]. The details are also found in the **Extended Experimental Methods** in **Supporting Information**.

In short, the biological samples included cultured human cancerous and non-cancerous cell lines spanning 9 different cancer types including brain (glioblastoma and neuroblastoma), blood (APL, AML, ALL), melanoma, BC, lung, HCC, CRC, OvC and BlaCa, **Supporting Table S1-S2**, and tumour and adjacent control tissue samples from six different cancer types including blood (CLL), non-melanoma skin (BCC and SCC) cancer, GC, HCC, CRC and PCa, **Supporting Table S3-S4**.

Proteins were extracted from the biological specimens, and the *N*-glycans released and handled as previously described [36, 37]. In short, the *N*-glycans were liberated using peptide:*N*-glycosidase F and quantitatively profiled in their reduced but otherwise non-derivatised alditol form using porous graphitised carbon liquid chromatography tandem mass spectrometry (PGC-LC-MS/MS) in negative ion polarity using linear and 3D ion trap mass spectrometers.

The raw data of all 467 LC-MS/MS glycomics datasets (available via the MassIVE Consortium, ftp://massive.ucsd.edu/MSV000083727/) were for the purpose of this study reinterrogated using the ESI-compass data analysis 4.0 software v1.1 (Bruker Daltonics) or Xcalibur v2.2 (Thermo Scientific) using assisting software i.e. GlycoMod (http://www.expasy.ch/tools/glycomod) and GlycoWorkBench v2.1 and manual *de novo* glycan sequencing as previously described [38, 39]. Glycan isomers were identified based on their monoisotopic precursor mass, MS/MS fragmentation pattern and their relative and absolute PGC-LC retention time [40]. The relative abundances of the individual *N*-glycans were determined from relative area-under-the-curve measurements based on extracted ion chromatograms performed for all charge states of the precursor ions using RawMeat v2.1 (Vast Scientific, www.vastscientific.com), QuantAnalysis software v2.1 (Bruker) and Skyline (64-bit) v20.1.0.76 [40]. Five major oligomannose species, M5-M9, some with two or more isobaric isomers were consistently identified across the investigated samples. Low levels of M4 and M9+Glc not fitting the common definition of oligomannose were observed in some samples and were for the sake of inclusion grouped with M5 and M9, respectively, for quantitation. **Supporting Data S2** (cell lines) and **Supporting Data S3** (tissues) summarise key information of all identified glycans including the oligomannosidic species for each cancer type i.e. details of their structure, composition, observed and theoretical molecular mass and mass deviation, PGC-LC retention, and relative abundance data.

### 2.3. Lectin flow cytometry

HL-60 cells (ATCC®, CCL-240™) were cultured in Roswell Park Memorial Institute-1640 media supplemented with 10% (v/v) heat-inactivated foetal bovine serum (Sigma-Aldrich) and maintained at 37°C with 5% CO2. Cells were pelleted by centrifugation at 300 x g, washed twice in phosphate buffered saline (PBS), and resuspended in PBS with 1% (w/v) bovine serum albumin (BSA, Sigma-Aldrich) before cell counting and viability determination using 0.4% (w/v) trypan blue staining (Sigma-Aldrich). A total of 5 x 10^5^ cells were used for each flow cytometry experiment. Cells were pelleted, washed once in PBS/1% BSA, resuspended in 100 µl concanavalin A-FITC (ConA-FITC, 1 µg/ml, Sigma-Aldrich) in the presence or absence of 0.5 M methyl-α-D-mannopyranoside (Me-α-Man) and then incubated in the dark on ice for 30 min. Cultured HL-60 cells without ConA-FITC treatment were used as a control. After incubation, cells were washed twice with PBS/1% BSA and resuspended in 500 µl PBS/1% BSA for analysis on a CytoFLEX S flow cytometer (Beckman Coulter, Australia). FITC was excited using a 488 nm laser and emission read at 525 nm using a 525/40 band-pass filter with a gain setting of 300. Data were acquired and stored in .fcs file format, imported into R v4.1.0 and analysed with the R-package flowCore v2.2.0.

### 2.4. Matrix-assisted laser desorption/ionization mass spectrometry imaging (MALDI-MSI) of tissue microarrays (TMAs)

TMA slides with 15 paired tumour and adjacent normal tissues from patients suffering from various cancer types were obtained (Biochain, USA, catalogue #T8235712-5, lot #B306118). The patient meta-data are provided in **Supporting Table S5**. The slides were pre-coated with indium tin oxide to generate a conductive surface for MALDI-MSI. TMAs were rehydrated and processed using a standard procedure for citric acid antigen retrieval [41]. Briefly, the tissue sections were heated at 60°C for 1 h on a heating block, washed twice in 100% (v/v) xylene for 5 min followed by serial dilutions in ethanol (100%, 90%, 70%, 50%, v/v) for 2 min each. Sections were then washed twice in Milli-Q water for 5 min. Next, the slides were boiled in 10 mM citraconic acid (pH 3.0) for 20 min on a steamer. Finally, tissue sections were immersed twice in 10 mM ammonium bicarbonate for 1 min and dried at room temperature in a humidity chamber. Peptide-*N*-glycosidase F (PNGase F PRIME™; N-Zyme Scientifics, USA) was sprayed onto the tissue sections using a TM-Sprayer (HTX-Imaging, USA) at a concentration of 0.1 µg/µl, flow rate of 25 µl/min, 15 passes, criss-cross pattern, velocity of 1200 mm/min, and track spacing of 3.0 mm. The nitrogen gas pressure was maintained at 10 psi. Tissue sections were incubated at 37°C for 2 h in a humidity chamber. A homogenous matrix (*α*-cyano-4-hydroxycinnamic acid, 10 mg/ml in 50% (v/v) acetonitrile / 0.1% (v/v) trifluoroacetic acid) was applied to tissues using the TM-sprayer. The settings were: 8 passes, 0.1 ml/min flow rate, 10 psi nitrogen pressure, 80°C capillary temperature, criss-cross pattern, velocity of 1200 mm/min, and track spacing of 3.0 mm.

The MS imaging data was acquired using a rapifleX MALDI-TOF/TOF mass spectrometer (Bruker Daltonics, Germany) controlled by flexControl (v4) and flexImaging (v5.1). Instrument settings were: *m/z* 920–3200, 5 kHz laser repetition rate and 2.5 GS/s. A total of 400 shots were acquired using a smartbeam 3D laser at 35 µm spatial resolution. The instrument was calibrated on-tissue using theoretical *m/z* values of known glycans prior to data acquisition. The MALDI-MSI data were analyzed using SCiLS Lab v2016b, SCiLS (Bruker Daltonics, Germany). Raw data were pre-processed by baseline subtraction and normalisation to the total ion count as previously shown [41, 42].

Following MALDI-MSI analysis, the matrix was eluted from the slides using 70% (v/v) ethanol and tissue sections were stained using hematoxylin and eosin to validate that investigated tissue spots were indeed from tumorigenic and non-tumour regions across the TMA using a standard protocol as described previously [41].

### 2.5. Mutation and expression data of human α1,2-mannosidases

Transcript (mRNA) expression data of MAN1A1, MAN1A2, MAN1B1 and MAN1C1 from select cancer cell lines were obtained using the RNAseq NCI-60 data [43] via the CellMiner^TM^ database (http://discover.nci.nih.gov/cellminer) [44]. Only cell lines from a primary tumour site and with matching glycomics data from whole cell lysates and/or microsomal fractions were included in this analysis. Thus, cancer cell lines from metastatic sites and with glycomics data of the secretome were left out for this purpose. Five cell lines matched these inclusion criteria i.e. HL-60 and CCRF-CEM (blood cancer), Hs-578T (BC), SKOV3 (OvC) and A549 (lung cancer).

Simple somatic mutation (SSM) and copy number variation (CNV) data of MAN1A1, MAN1A2 and MAN1B1 (which were found to dictate the oligomannose expression levels in cancer cells) were obtained from The Cancer Genome Atlas (TCGA) (https://www.cancer.gov/tcga) [45]. We only considered data obtained from unique cancer primary sites including 11 projects with the following accession IDs (primary site): TCGA-SKCM (skin), TCGA-UCEC (uterus), TCGA-COAD (colon), TCGA-LUSC (lung), TCGA-STAD (stomach/gastric), TCGA-BLCA (bladder), TCGA-OV (ovary), TCGA-BRCA (breast), TCGA-LIHC (liver), TCGA-KIRC (kidney) and TCGA-PRAD (prostate). The mean loss/gain ratio of the SSMs was determined by calculating the average frequency (cases affected relative to cases tested) of CNV loss events divided by CNV gain events across the 11 cancer projects.

Transcript (mRNA) expression data of MAN1A1 (which showed a high rate of deleterious mutations across cancers) from paired and unpaired tumour and non-tumour tissues across 10 cancer types were obtained from the Gene Expression Omnibus (GEO) database (https://www.ncbi.nlm.nih.gov/geo/) [46, 47]. Projects with the following GEO accession IDs (primary site) were considered: GDS1375 (skin), GDS3975 (uterus), GDS4382 (colon), GD3837 (lung), GDS1210 (gastric), GDS1479 (bladder), GDS2835 (ovary), GDS3139 (breast), GDS505 (kidney) and GDS2545 (prostate). Only human cancer types covered by our literature survey and/or our collection of glycomics datasets were included in this analysis. Other selection criteria included the availability of data from both non-tumour and tumour (primary site) tissues of a minimum of five patients per cohort.

### 2.6. Statistics

Statistical tests of the glycomics data and the α1,2-mannosidase transcript data were performed using paired and unpaired one- or two-tailed Student’s t-tests. Significance has been consistently indicated as * (*p* < 0.05), ** (*p* < 0.001), *** (*p* < 0.0001) while n.s. denotes a lack of significance (*p* ≥ 0.05). The number of patient samples (n) for the different tissue analyses has been indicated. Cell line glycoprofiling data were generated from at least two or three technical replicates for each experiment. Data have generally been plotted as the mean ± standard deviation (SD) where n ≥ 3.

## 3. Results & Discussion

### 3.1. Literature suggests wide-spread associations between oligomannose and cancer

Despite forming a large and vital part of the human *N*-glycome, the role(s) of oligomannosylation in cancer remains under-studied. To this end, we firstly performed a comprehensive and unbiased survey of the original research literature published over the past two decades in the field of glyco-oncology to test for a possible link between oligomannosylation and human cancers. Interestingly, we identified a considerable body of research papers that consistently reported on oligomannose elevation across 10 cancer types including bladder (BlaCa) [48], breast (BC) [49], cholangiocarcinoma (CC) [50], colorectal (CRC) [33], endometrial [51], liposarcoma [52], lung [53], ovarian (OvC) [54], oral cancer (OC) [55], and prostate cancer (PCa) [56], **Table 1**. These scattered but consistent observations reported by various research groups around the world were made from a diverse set of cancer specimens (cell lines, tissues and bodily fluids) using different analytical techniques (MS, HPLC, and lectin arrays). Importantly, the cancer-associated elevation of oligomannose was supported by multiple independent studies for several cancer types. Notably, we also found papers describing unchanged or reduced levels of oligomannosylation in five other cancer types including gastric (GC) [57], hepatocellular carcinoma (HCC) [58], kidney (KC) [59], pancreas (PanCa) [60], and thyroid cancer (TC) [61]. Thus, our initial literature survey pointed to a strong association between oligomannosidic *N*-glycans and a subset of human cancers.

### 3.2. Re-interrogation of pan-cancer LC-MS/MS glycomics datasets

Guided by the literature survey, we then revisited our collection of 467 PGC-LC-MS/MS glycomics datasets previously compiled for a recent pan-cancer study [37] to systematically test for an association between oligomannosylation and cancer. Our quantitative glycome datasets covered 11 types of human cancers including brain (glio- and neuroblastoma), BlaCa, blood (APL, AML, ALL), skin (melanoma and non-melanoma, BCC and SCC), BC, lung, HCC, GC, CRC, PCa and OvC spanning both an extensive set of human cell lines and cohorts of paired and unpaired tissue samples. Importantly, all *N*-glycome datasets were acquired using the same analytical technique enabling not only relative *N*-glycan quantitation within each sample, but also accurate comparisons within and between sample cohorts.

The five common oligomannosidic *N*-glycans, M5-M9 (depicted in Figure 1), were consistently identified and quantified across all samples. As expected from the biosynthetic trimming process, isomers of all five oligomannosidic *N*-glycans were identified across most samples. The less common M4 and M9+Glc were observed at low abundance in some samples. The spectral evidence can be found in **Supporting Data S1** and all *N*-glycome profile data are tabulated in **Supporting Data S2-S3**.

### 3.3. Pan-cancer cell line glycoprofiling demonstrates that oligomannose is a cancer-wide signature

Our glycomics data of 34 human cell lines spanning 9 cancer types showed that oligomannosidic *N*-glycans are widely and abundantly expressed in cancerous cells, **Supporting Table S1-S2**. While the total oligomannose levels varied considerably across the investigated cancer types and protein fractions (5.3%-91.2% of the *N*-glycome), the whole cell lysate and membrane fractions were particularly rich in oligomannose (>20%), **Figure 2A** and **Supporting Table S6**. In line with our previous observation [62], less oligomannosylation was consistently found in the secretome of cultured cancer cells indicating that oligomannosidic *N*-glycans are mainly features of the cellular component of cancer cells. Incompletely processed glycoproteins still trafficking the secretory pathway may contribute significantly to the high levels of oligomannosylation within cancer cells. Despite their relatively low secretion rate from cancers cells, oligomannosidic *N*-glycans have repeatedly been found to be elevated in the blood of cancer patients including those suffering from BC, CC and lung cancers [63–65] relative to the low oligomannose levels found in the blood of healthy individuals [66, 67].

**Figure 2.**
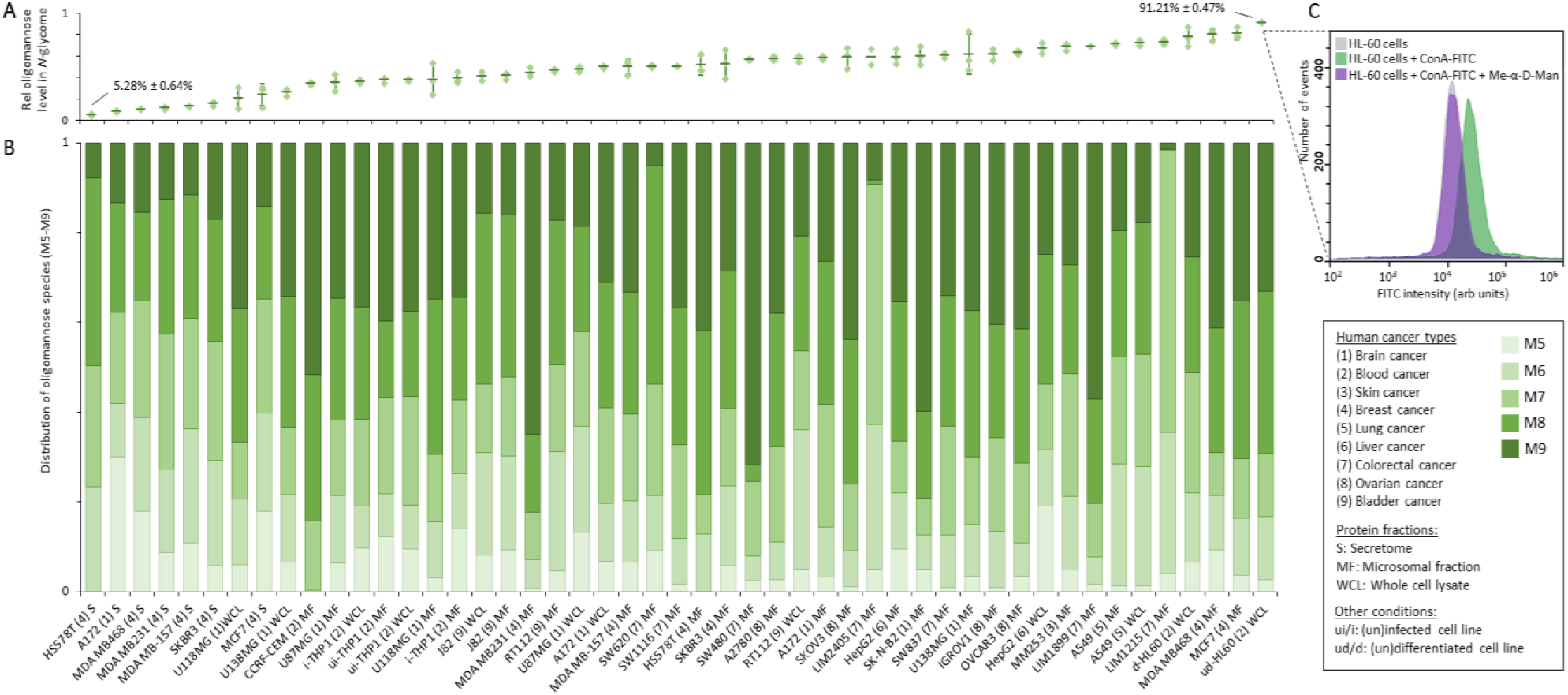
Oligomannosylation is a prominent feature across human cancer cell lines. (**A**) Total oligomannose level and (**B**) the relative distribution of individual oligomannosidic species (M5-M9) of the *N*-glycome from different protein fractions (secretome, microsomal fractions, whole cell lysates) of all investigated 34 human cancer cell lines spanning nine cancer types. See **Supporting Table S1-S2** for details of the studied cell lines. For A and B, data were either plotted as the mean ± SD (for n ≥ 3, technical replicates) or as the mean (for n = 2, technical duplicates). See **Supporting Data S1** for spectral evidence and **Supporting Data S2** and **Supporting Table S6-S7** for raw and tabulated quantitative glycome data, respectively. (**C**) Cell surface expression of oligomannosidic epitopes of the oligomannose-rich HL-60 cell line was shown by the considerable surface reactivity of the mannose-recognising concanavalin A (ConA, FITC-conjugated, green trace) using lectin flow cytometry. The ConA reactivity could be competitively inhibited by methyl-α-D-mannopyranoside (Me-α-D-Man) (purple trace) bringing the signal to background HL-60 levels without ConA-FITC treatment (grey trace).

The M7-M9 glycans were the most common oligomannosidic species expressed by the cancer cell lines pointing to a significant glycan under-processing in cancer, **Figure 2B** and **Supporting Table S7**. Glycan processing may be impacted by several cellular factors e.g. rate of protein synthesis, cellular growth rate (doubling time), protein trafficking time, and the levels of multiple glycan-processing enzymes and nucleotide sugars [68] as well as various protein factors [69]. While the reduced expression of several α1,2-mannosidases catalysing the M9-to-M5 trimming process has been reported in cancer (discussed below) [70–72], the possible contribution of these many other cellular and protein factors to the oligomannose-rich glycophenotypes of cancer cells remains unexplored.

Comparative glycoprofiling of a few “normal” (non-tumourigenic) immortalised cell lines derived from breast (HMEC) and ovarian (HOSE 6.3 and HOSE 17.1) tissues and their matching cancerous cell lines from the same tissue origins including Hs-578T, MCF-7, MDA-MB-157/231/468 and SK-BR-3 breast cancer cell lines and A2780, IGROV1 and SKOV3 ovarian cancer cell lines indicated an elevation of oligomannose in the cancer cell lines (data not shown), a relationship explored in greater details below using paired tissue samples.

Cell surface glycoepitopes are known to play important roles in cell-cell and cell-extracellular matrix communication [1, 24, 73]. To this end, we explored the cell surface expression of oligomannosidic epitopes of the promyelocytic HL-60 cell line using lectin flow cytometry. Significant surface reactivity to the mannose-recognising lectin, concanavalin A (ConA, FITC-conjugated) was observed, **Figure 2C** (green trace). Importantly, the ConA reactivity could be competitively inhibited by methyl-α-D-mannopyranoside (Me-α-D-Man) (purple trace) bringing the signal back to the base levels established for HL-60 without ConA-FITC treatment (grey trace). The cell surface expression of oligomannosidic epitopes on HL-60 cells and possibly other cancer cells is interesting as it suggests roles of oligomannose in cancer cell communication and may open for novel glycan-centric diagnostic approaches as well as new treatment strategies such as chimeric antigen receptor T-cell therapy directed to surface-exposed oligomannose [74].

### 3.4. Comparative tissue glycomics and MALDI-MS imaging confirm oligomannose elevation in a subset of cancer types

Next, *N*-glycomics data from 126 paired and unpaired tumour and non-tumour tissues (fresh frozen, FF and formalin-fixed paraffin-embedded, FFPE) from various patient cohorts spanning seven cancer types were re-analysed, **Supporting Table S3-S4.** A subset of the investigated cancer types including non-melanoma BCC (FF, n = 14; FFPE, n = 20) and SCC (FFPE, n = 15), and CRC (FF, n = 5-6) showed strong tumour-associated elevation of oligomannose (*p* < 0.0001 for all three cancer types, paired two-tailed t-tests), **Figure 3A**. Most of the individual oligomannosidic *N*-glycan species showed a significant elevation in the tumour tissues relative to the adjacent control tissues albeit no consistent patterns of the relative M5-M9 distribution between cohorts and cancer types could be identified, **Figure 3B-D**.

**Figure 3.**
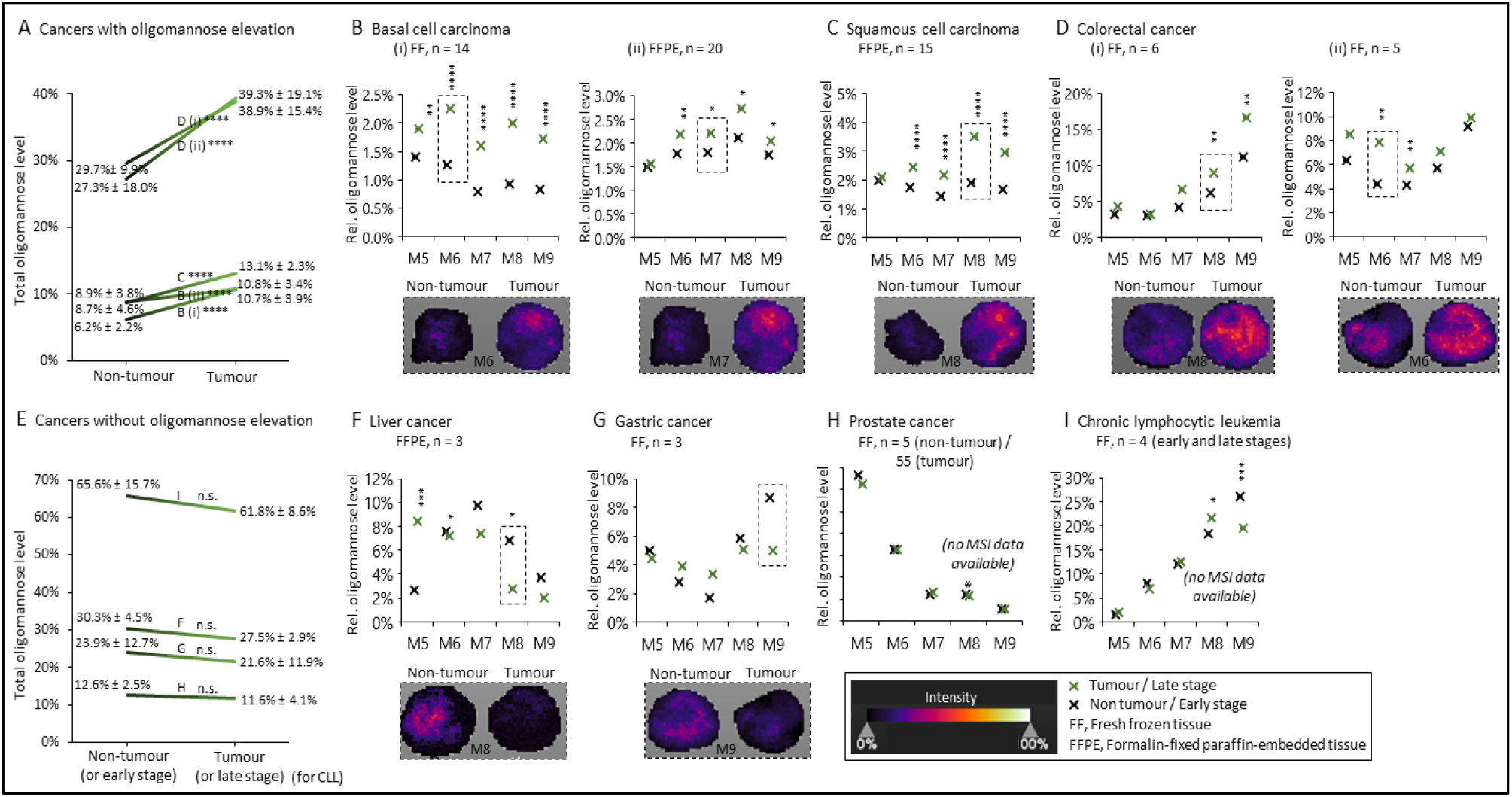
Tissue glycomics and MALDI-MSI data show oligomannose elevation in a subset of cancers. (**A**) Some cancer types displayed elevated oligomannose levels as demonstrated by quantitative glycomics of donor-paired tumour and non-tumour tissues from patients with (**B**) basal cell carcinoma (i, FF; ii, FFPE tissues), (**C**) squamous cell carcinoma and (**D**) colorectal cancer (i-ii, FF tissues). See **Supporting Table S3-S4** for details of the investigated tissues. (**E**) Another subset of cancers did not exhibit oligomannose elevation as assessed via quantitative glycomics of paired tumour and non-tumour tissues from patients with (**F**) liver cancer and (**G**) gastric cancer and unpaired tumour and non-tumour tissues from patients with (**H**) prostate cancer and patients with (**I**) early- or late-stage chronic lymphocytic leukemia. See **Supporting Data S3** and **Supporting Table S8-S9** for all raw and tabulated glycome data, respectively. Statistics was performed using paired and unpaired two-tailed t-tests (n, patient samples, as indicated). \**p* < 0.05, \*\**p* < 0.001, \*\*\**p* < 0.0001, n.s. not significant (*p* ≥ 0.05). The MALDI-MSI data were from TMAs of unpaired tumour and non-tumour tissues. Examples of oligomannosidic *N*-glycans displaying prominent regulation in tumours are provided (boxed, broken line). No MSI data could be obtained for prostate cancer and chronic lymphocytic leukemia. See key for MALDI-MSI intensity range and **Supporting Table S5** for details of the TMAs used for the MALDI-MSI.

MALDI-MSI of glycans, a powerful tool to spatially profile liberated glycans from biological specimens including TMAs (as used here, see **Supporting Table S5**) [42, 75], recapitulated the glycomics data by consistently showing a higher expression of specific oligomannosidic *N*-glycans in the tumour relative to non-tumourigenic tissues from matching cancer types, **Figure 3B-D inserts**. Hematoxylin and eosin staining confirmed that the investigated spots were indeed from tumour and non-tumour regions across the TMA (data not shown). While this study did not address the cellular origin(s) of the elevated oligomannose in the heterogeneous tumour micro-environment, MALDI-MSI and glycoproteomics with cell annotation of ovarian [54] and prostate [75, 76] tumour tissues have previously indicated that oligomannose-rich glycophenotypes map to cancer cells as oppose to the stroma, immune cells and other cell types forming important components of the tumour microenvironment. Collectively, these studies therefore indicate that raised oligomannose levels originate directly from the cancer cells, however, this aspect and the still elusive protein carriers of oligomannose require further investigation across cancer types. Such studies can now be carried out with emerging structure-focused glycoproteomics methods [77] and advanced cell-sorting and characterisation technologies [78].

In keeping with observations captured by our literature survey, several cancer types showed unchanged or slightly reduced levels of oligomannose in tumour tissues including HCC (FFPE, n = 3) and GC (FF, n = 3) (*p* ≥ 0.05 for all cancer types, paired two-tailed t-tests) and PCa (FF, n = 5 and 55) (*p* ≥ 0.05, unpaired two-tailed t-tests) and the stage-wise progression of CLL (FF, n = 4) (*p* ≥ 0.05, unpaired two-tailed t-tests), **Figure 3E**. While late-stage CLL did not show oligomannose elevation relative to early-stage disease, these samples notably all displayed very high levels of oligomannosylation (>60%). Our data therefore did not point to a progression-dependent elevation of oligomannose in CLL, but follow-up analysis of B cells from healthy donors are still required to determine if raised oligomannose is a disease characteristic of CLL.

Generally, M5-M6 were prominent glycans expressed in the subset of cancer types that did not display oligomannose elevation albeit variable patterns were observed between cancers, **Figure 3F-I** and **Supporting Table S8-S9**. Reduced oligomannose levels in HCC and GC tumour tissues were observed using MALDI-MSI supporting the down-regulation of specific oligomannose species in these cancer types, **Figure 3F-G inserts**. The tumour tissues from HCC and GC instead expressed high levels of complex-type *N*-glycans suggesting a higher degree of glycan processing and consequently a greater influence of other glycan-processing enzymes (e.g. fucosyl-, GlcNAc-, galactosyl- and sialyl-transferases) in those cancers.

### 3.5. Three α1,2-mannosidases dictate oligomannose levels in cancer

We then investigated a possible connection between the biosynthetically-relevant α1,2-mannosidases and oligomannose expression in cancer cells by firstly retrieving transcript (mRNA) data of MAN1A1, MAN1A2, MAN1B1 and MAN1C1 in five cancer cell lines (HL-60, CCRF-CEM, Hs-578T, SKOV3 and A549) from the Cell Miner^TM^ database, **Figure 4A**. Correlation analyses of the transcript data and our glycomics data of the same cell lines showed that oligomannose levels correlated negatively with the expression of MAN1A1 (r = - 0.809), MAN1A2 (r = -0.614) and MAN1B1 (r = -0.966) while MAN1C1 showed a relatively weak positive correlation (r = +0.728). Thus, this analysis indicated that reduced MAN1A1, MAN1A2, and MAN1B1 expression promotes an oligomannose-rich glycophenotype in cancer, an intuitive relationship given the known involvement of these α1,2-mannosidases in the M9-to-M5 glycan trimming process (see Figure 1). While the relative contribution of MAN1C1 (and MAN1A1/A2/B1) to the oligomannose trimming remains ill defined, recent studies have suggested that MAN1C1 plays unexpected anti-cancer roles in tumour suppression [71, 79], perhaps explaining its opposite expression pattern relative to the other α1,2-mannosidases linked to pro-tumorigenic processes (discussed below) [70, 79, 80].

**Figure 4.**
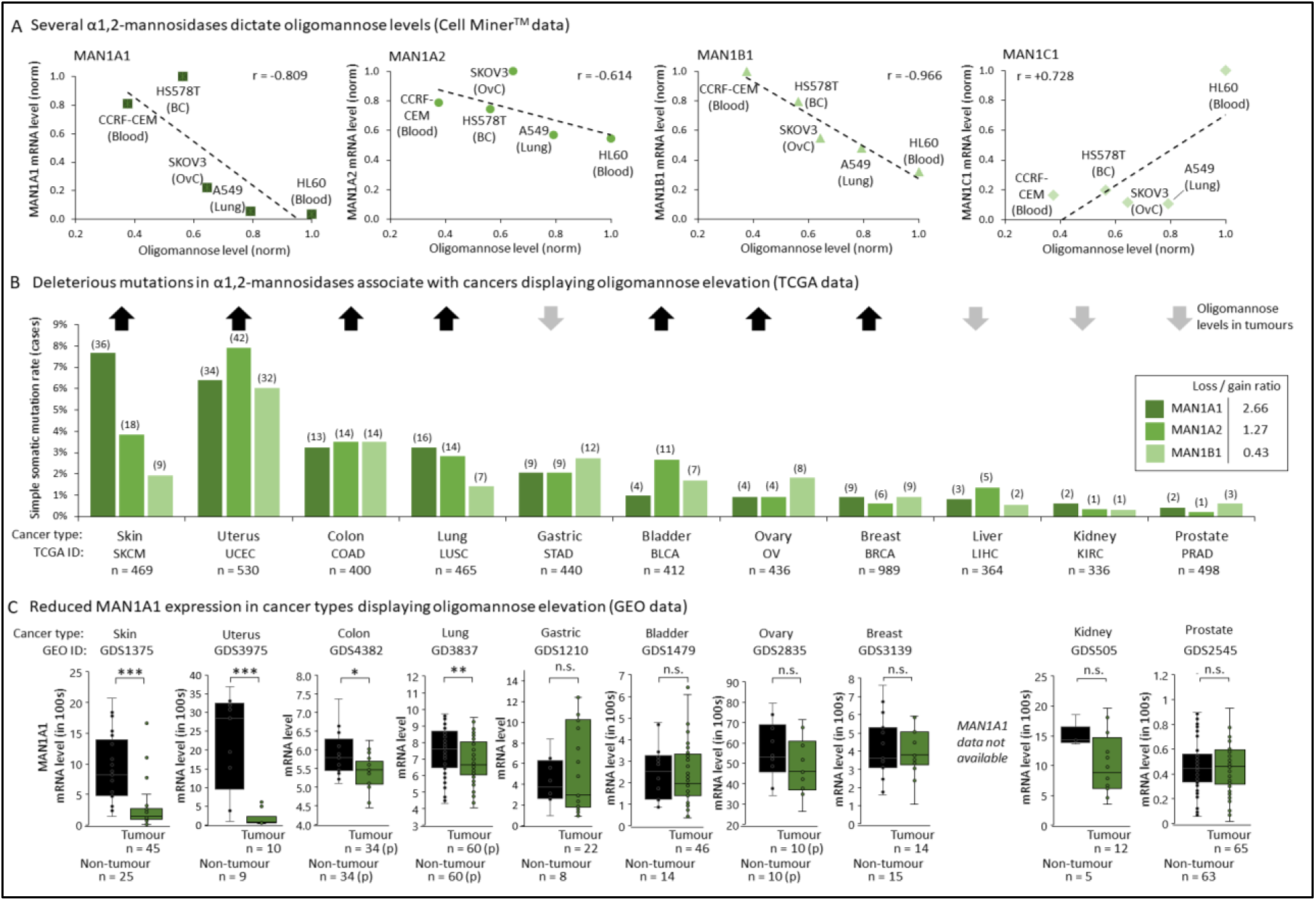
α1,2-mannosidase-mediated regulation of oligomannose expression in cancer. (**A**) Correlation analysis between the expression (mRNA) of biosynthetically-relevant α1,2-mannosidases (MAN1A1, MAN1A2, MAN1B1, and MAN1C1 [2], see Figure 1), as extracted from the Cell Miner^TM^ database, and the total oligomannose levels in five cancer cell lines covered by both our glycomics data (Figure 2) and the Cell Miner^TM^ database. Notably, MAN1A1, MAN1A2, and MAN1B1 showed a negative correlation with the oligomannose levels. (**B**) Frequency of simple somatic mutations (SSMs) of MAN1A1, MAN1A2 and MAN1B1 determined based on data retrieved from The Cancer Genome Atlas (TCGA) across 11 cancer types investigated in this study. Insert: Analysis of the gene copy number variation (CNV) loss- or gain-of-function ratio of each α-mannosidase across the 11 cancer types using TCGA data showed that deleterious mutations in MAN1A1 is a prominent cancer feature. The up- (black arrows) and down-regulation/unchanged levels (grey arrows) of oligomannose in tumours based on literature (Table 1) and/or our glycomics data (Figure 3) is indicated for each cancer type to aid the data interpretation. (**C**) MAN1A1 mRNA expression in paired (p) and unpaired tumour and non-tumour tissues across cancer types based on data extracted from the Gene Expression Omnibus (GEO). Reduced MAN1A1 expression was observed for cancers displaying oligomannose elevation. Statistics was performed using paired and unpaired two-tailed t-tests (n, sample numbers, as indicated). \**p* < 0.05, \*\**p* < 0.001, \*\*\**p* < 0.0001, n.s. not significant (*p* ≥ 0.05).

### 3.6. Deleterious mutations and low expression of MAN1A1 in oligomannose-rich cancers

Next, we determined the frequency of simple somatic mutations (SSMs) of MAN1A1, MAN1A2 and MAN1B1 (dictating the levels of oligomannose in cancer, see above) across 11 cancer types (skin, uterus, colon, lung, gastric, bladder, ovary, breast, liver, kidney and prostate) using data retrieved from The Cancer Genome Atlas (TCGA), **Figure 4B**. To enable comparisons across datasets, the cancer types were selected to match the cancers covered by our glycomics data and the literature survey. Interestingly, relatively high mutation rates of all three α1,2-mannosidases were found in cancer types displaying oligomannose elevation i.e. skin, uterus, colon, and lung cancers while lower α1,2-mannosidase mutation rates were observed in liver, kidney and prostate cancers not featuring elevated oligomannosylation. Moreover, analysis of the gene copy number variation (CNV) as a proxy for a loss- or gain-of-function of the SSMs indicated that the raised oligomannose levels in the investigated cancer types may arise predominantly from deleterious mutations of MAN1A1 (loss/gain ratio = 2.66) as oppose to MAN1A2 and MAN1B1 (loss/gain ratio = 1.27 and 0.43, respectively), **Figure 4B insert**.

The MAN1A1 transcript expression was therefore investigated in the same cancer types using Gene Expression Omnibus (GEO) data collected from paired and unpaired tumour and non-tumour tissues, **Figure 4C**. In line with trends observed from the mutation data, reduced MAN1A1 expression (all *p* < 0.05) was observed in tumour tissues belonging to the oligomannose-rich cancers including skin, uterus, colon and lung cancer, while cancer types not showing oligomannose elevation (gastric, kidney, and prostate) showed unchanged MAN1A1 expression (all *p* ≥ 0.05).

Supporting our MAN1A1-centric findings, MAN1A1 has repeatedly been reported to be down-regulated in a variety of metastatic cancer cell lines including in HCC [81], CC [50], OvC [80] and PCa [82] relative to their non-metastatic counterparts. Thus, MAN1A1 and/or the resulting oligomannosidic *N*-glycan products may be involved (directly or indirectly) in the regulation of the growth and dissemination of some cancer types. Moreover, a recent study showed that reduced MAN1A1 expression and oligomannose elevation are features of advanced-stage BC tumour tissues with high tumour aggressiveness relative to early-stage BC tumour tissues and that these characteristics lead to impaired survival in BC patients [70].

We found comparably less literature linking the other biosynthetically relevant α1,2-mannosidases to cancer. In a recent study, MAN1B1 expression was reported to be up-regulated in BlaCa relative to normal tissues and found to be a regulator of cell proliferation [72]. As mentioned above, other studies have indicated that MAN1C1 is a tumour suppressor in KC [71] and HCC [79]. Finally, low expression of MAN1C1 (and a concomitant up-regulation of MAN1A1) was also reported in HCC (hepatitis B-positive) stage 1 relative to stage 2 and 3 further supporting that MAN1C1 plays roles in tumour suppression, whereas MAN1A1 was suggested as a novel oncogene in hepato-carcinogenesis [79]. We did not find any literature reporting on the role of MAN1A2 in cancer.

Collectively, our glycomics data, the existing glyco-oncology literature and data retrieved from public cancer repositories provide robust evidence suggesting that MAN1A1 is a key α1,2-mannosidase that dictates oligomannose expression in a subset of human cancers.

## 4. Conclusions

Guided by a comprehensive literature survey pointing to a hitherto under-explored association between oligomannosylation and human cancers, we have here re-interrogated a large volume of PGC-LC-MS/MS glycomics datasets generated from a diverse set of cancer cell lines and valuable cohorts of patient tissues. Our quantitative glycomics data and supporting MALDI-MSI data clearly demonstrate that raised oligomannosylation is a strong molecular feature of a subset of human cancer types. Notably, lectin flow cytometry data indicated that oligomannose is a prominent cancer cell surface epitopes. Finally, data from well-curated cancer repositories suggested that deleterious mutations and reduced expression of key α1,2-mannosidases, particularly MAN1A1, are responsible for the oligomannose elevation observed in a subset of human cancers. Collectively, these findings open hitherto unexplored avenues for the development of new enzyme- and glycan-directed cancer biomarkers and therapeutic targets.

## Supporting information

Extended Experimental Methods and Supporting Tables S1-S9

## Abbreviations

2-AB: 2-aminobenzoic acid
ALL: acute lymphocytic leukaemia
AML: acute monocytic leukaemia
APL: acute promyelocytic leukaemia
APTS: 8-amino-1,3,6-pyrenetrisulfonic acid
Asn: asparagine
BC: breast cancer
BCC: basal cell carcinoma
BlaCa: bladder cancer
BPH: benign prostatic hyperplasia
BSA: bovine serum albumin
CC: cholangiocarcinoma
CLL: chronic lymphocyte leukaemia
CNV: copy number variation
ConA: concanavalin A
CRC: colorectal cancer
ER: endoplasmic reticulum
FF: fresh frozen
FFPE: formalin-fixed paraffin-embedded
Fuc: fucose
Gal: galactose
GC: gastric cancer
GEO: Gene Expression Omnibus
Glc: glucose
GlcNAc: *N*-acetylglucosamine
HCC: hepatocellular carcinoma
KC: kidney cancer
LC-MS/MS: liquid chromatography tandem mass spectrometry
MALDI-MSI: matrix-assisted laser desorption / ionization mass spectrometry imaging
Man: mannose
Me-α-Man: methyl-α-D-mannopyranoside
NeuAc: *N*-acetylneuraminic acid
OC: oral cancer
OvC: ovarian cancer
PanCa: pancreas cancer
PBS: phosphate buffered saline
PCa: prostate cancer
PGC: porous graphitised carbon
SCC: squamous cell carcinoma
SD: standard deviation
SSM: simple somatic mutation
TC: thyroid cancer
TCGA: The Cancer Genome Atlas
TMA: tissue microarray

## Conflicts of Interest

The authors have declared no conflict of interest.

## Author Contributions

Conceptualisation, S.C. and M.T.A.; methodology, S.C., J.U. and A.E.D; software, S.C., R.K. and A.E.D; validation, S.C. and M.T.A.; formal analysis, S.C., R.K. and J.U.; investigation, S.C., R.K., and M.T.A.; resources, A.E.D. and M.T.A.; data curation, S.C. and M.T.A.; writing— original draft preparation, S.C. and M.T.A.; writing—review and editing, S.C., R.K., J.U., L.Y.L., A.E.D. and M.T.A.; supervision, M.T.A.; project administration, M.T.A.; funding acquisition, M.T.A. All authors have read and agreed to the published version of the manuscript.

## Acknowledgements

S.C. was supported by an International Macquarie Research Excellence Scholarship Program. R.K. was supported by an Early Career Fellowship from the Cancer Institute New South Wales. J.U. was supported by a Macquarie Research Excellence Scholarship Program. A.E.D. was supported by the Griffith University Institute for Glycomics. M.T.A. was supported by a Macquarie University Safety Net Grant. The authors thank Russel M. Vincent for valuable assistance with the acquisition and analysis of flow cytometry data.

## Notes

### Competing Interest Statement

The authors have declared no competing interest.

