## Extended Experimental Methods and Supporting Tables S1-S9 for "Oligomannosylation and MAN1A1 expression associate strongly with a subset of human cancer types"

for

### Table of Content of the SI


| | |
|---|---|
| Extended Experimental Methods | Page 3-13 |
| Supporting Tables | |
| Supporting Table S1 | Page 14-17 |
| Supporting Table S2 | Page 18-21 |
| Supporting Table S3 | Page 22-23 |
| Supporting Table S4 | Page 24-29 |
| Supporting Table S5 | Page 30-30 |
| Supporting Table S6 | Page 31-32 |
| Supporting Table S7 | Page 33-34 |
| Supporting Table S8 | Page 35-35 |
| Supporting Table S9 | Page 36-36 |
| References | Page 37-40 |

### Extended Experimental Methods

#### Human ethics

Ethics approvals were obtained from the local ethic committees relevant to the individual sample cohorts investigated in this study. Specifically, the study of i) chronic lymphocyte leukaemia (CLL) tissues were approved by the human research ethics committee (HREC) at the University of Sydney, Australia (Project #8935), ii) non-melanoma tissues (basal cell carcinoma, BCC and squamous cell carcinoma, SCC) were approved by ethics committee at University of Leipzig (ethics #127-11-18042011) iii) gastric tissues were approved by *Comitato Etico Regione Toscana* (Tuscany, Italy) and confirmed to the ethical guidelines of the Declaration of Helsinki (1975), iv) cirrhotic and hepatocellular carcinoma (HCC) liver tissues were approved by the ethics committee at *Klinička Bolnica Merkur* at Zagreb, Croatia v) colorectal cancer (CRC) tissues were obtained from Sydney South West Area Health Service (ethics #X08-0614) and Department of Pathology, Yonsei University and Liver Cancer Specimen Bank of the National Research Resource Bank program of the Korea Science and Engineering Foundation of the Ministry of Science and Technology (Seoul, South Korea) (ethics #5201100040) respectively, vi) prostate cancer (PCa) and matching benign prostatic hyperplasia (BPH) tissues were approved by Faculdade de Medicina do Estado de São Paulo (Protocol nº2695126), vii) ovarian cancer tissues were approved by HREC at Royal Adelaide Hospital, Adelaide, Australia (#140101). Finally, adipose tissue samples from healthy donors were collected as controls after approval from the HREC at Macquarie University, Sydney, Australia (#5201100385). The appropriate biosafety measures were followed while isolating and handling the human cancer cell lines and tissues.

### Chemical and reagents

Ultra-pure water was sourced from a Milli-Q system (Merck/Millipore). Recombinant *Escherichia coli* produced peptide-*N*-glycosidase F (PNGase F) from *Flavobacterium meningosepticum* and *Elizabethkingia miricola* were from Roche Diagnostics (Mannheim, Germany) and Promega (Sydney, Australia), respectively. Other chemicals, reagents and proteins were obtained from Sigma Aldrich or Thermo Scientific unless specified. Dulbecco's Modified Eagle's medium (DMEM), MCDB 105: Medium 199 (1:1, v/v), Minimum Essential Medium (MEM alpha), Roswell Park Memorial Institute medium 1640 (RPMI), and foetal bovine serum (FBS) were from Sigma Aldrich or Thermo Fisher Scientific (Sydney, Australia).

### Human cancer cell lines

Human cell lines were obtained from the American Type Culture Collection (ATCC) (Manassas, VA), Cell Bank Australia (CBA) (Sydney, Australia), Cell Line Services (CLS) (Eppelheim, Germany), Garvan Institute of Medical Research (Sydney, Australia) and from multiple other sources as indicated in **Supporting Table S1-S2**. These cell lines were collected and studied over many years by many past and present members and collaborators of our glycoanalytical laboratories. The cell lines were generally stored at -140°C in 90% (v/v) FBS and 10% (v/v) dimethyl sulfoxide (DMSO) until use. All cultured cells were confirmed mycoplasma-free prior to the *N*-glycome profiling.

### Tumour and non-tumour tissues from cancer patient cohorts

Tissue samples were obtained from multiple cancer patient cohorts or from various control individuals suffering from other non-cancer conditions at a range of hospitals and clinics as described in **Supporting Table S3-S4**. Many of these tissue samples have been described in recent publications [1-10]. In brief, tumourigenic and non-tumourigenic tissues were obtained by surgery by trained clinicians after obtaining written and informed patient consent, see above and **Supporting Table S3** for ethics details.

**Cell culture conditions**

The human cancer cell lines were thawed and cultured in DMEM, RPMI 1640, MEM or MCDB 105 media in Corning cell culture flasks (see **Table 1A** for an overview of culture conditions). These culture media differed slightly in their nutritional composition, but all contained essential components including glucose, L-glutamine, sodium pyruvate, sodium bicarbonate, phenol red as pH indicator and salts. The media were supplemented with 10% (v/v) FBS, 1% (v/v) penicillin and 100 U/ml streptomycin, and, in some cases, 25 U/l insulin, 1 µM α-thioglycerol, 1 mg/ml hydrocortisone and non-essential amino acids. FBS-free media was used to obtain the secretome of some cultured cells (see below). All cells were grown at 37°C, 5% $CO_2$. The cell viabilities were regularly determined using trypan blue exclusion, counted using an automated cell counter (BioRad) and visualised using an inverted light microscope (Olympus) at 10-40x magnification to monitor the morphology of the cells. The cells were sub-cultured at 90-95% confluency typically every 72-96 h [11].

<u>Sub-culturing of non-adherent cancer cells:</u> Suspension cells were collected by centrifugation of the culture media at 500 x g for 10 min

at 20°C. The cell pellet fractions were re-suspended in culture media and transferred to new flasks. Fresh media was added every 48 h.

Sub-culturing of adherent cancer cells: The culture media were removed, and the cells were washed with sterile-filtered phosphate buffered saline (PBS) and quickly detached by a brief treatment with 0.25% (w/v) trypsin at 37°C. Fresh media was added (1:1, v/v) to quench the activity of trypsin and the cells were transferred to four new flasks. Fresh media were added every 48 h.
For all cultured cells, the cell pellets were thoroughly washed and re-suspended in cold PBS to remove traces of FBS and then centrifuged prior to storage at -30°C until being used for analysis.

**Tissue sample handling**

The below describes the sample handling of the tissues obtained from patients suffering from different cancers investigated in this study.

Chronic lymphocytic leukaemia (CLL): Frozen CLL samples (~20 x $10^6$ mononuclear cells) were thawed in a 37°C water bath, mixed with MACS buffer in a polypropylene round-bottom tube and centrifuged at 400 x g for 5 min at 20°C. The resulting cell pellets were re-suspended in MACS buffer, magnetically isolated using CD19-conjugated microbeads and incubated at room temperature for 15 min. The samples were again centrifuged, cell pellets were re-suspended in MACS buffer, and the B cells were isolated on MACS LS columns. The column was washed with MACS buffer to remove any unbound cells and then removed from the magnet and placed on a 15 ml CellStar tube. MACS buffer was loaded into the column and flushed out immediately by firmly pushing the plunger into the column to expel the liquid containing cells with microbeads. The process was repeating several times. The tubes were centrifuged as described above. The resulting cell pellets were lysed using probe sonication (5 s cycles, 20% amplitude) on ice and proteins were extracted by chloroform-water-methanol extraction (see below).

Basal cell carcinoma (BCC) and squamous cell carcinoma (SCC): To allow for an accurate dermato-pathological investigation, only patients exhibiting a well-circumscribed BCC or SCC of at least 5 mm diameter were included in the study. The BCC and SCC tissues were obtained by fusiform excision with micrographic control of the margins under local anaesthesia. Punch biopsies were obtained from the tumourigenic and normal regions of the excised tissues, snap-frozen in liquid nitrogen and stored at -80°C. Prior to use, the samples were washed three times with 70% (v/v) ethanol and then three times with 50 mM ammonium bicarbonate. The samples were homogenised in a lysis buffer containing 50 mM ammonium bicarbonate, 1 M urea, 10% (v/v) acetonitrile (ACN) and 0.1% (w/v) sodium dodecyl sulfate (SDS) using a IKA T10 Ultraturrax homogeniser (Staufen, Germany) and then sonicated using a tabletop Branson sonifier B-12 sonicator for 30 s. To increase the protein/peptide solubility, 4 µg trypsin was added, and the samples were incubated overnight at 37°C before insoluble material/particles were removed by centrifugation at 14,000 x g for 30 min. The supernatant fractions were collected and the protein complement fractions were isolated by chloroform-water-methanol extraction (see below).

Hepatocellular carcinoma (HCC): Fresh liver tumour tissues were micro-dissected and prepared for hematoxylin-eosin staining to enable a tissue sampling containing at least 70% tumour cell content [12] before snap-frozen in liquid nitrogen and stored at -80°C. Prior

to use, the frozen tissue blocks were homogenised on ice using an IKA T10 Ultraturrax homogeniser (Staufen, Germany) in lysis buffer containing 50 mM Tris-HCl (pH 7.4), 100 mM NaCl, and 1 mM ethylenediaminetetraacetic acid (EDTA) with 5 mM EDTA-free protease inhibitor tablets. The solutions were then sonicated on ice using a tabletop Branson sonifier B-12 sonicator for 10-30 s and centrifuged at 2,000 x g for 20 min at room temperature. The supernatant fractions were collected and the proteins isolated using microsomal preparation (see below) [13].
The formalin-fixed paraffin-embedded (FFPE) tissue samples from BCC, SCC and liver were treated with 10% (v/v) formalin for 24 h followed by washing with 70%, 95% and 100% (all v/v) ethanol (2 x 1 h each), xylene (2 x 1 h at 37°C) and then liquid paraffin (3 x 1 h at 60°C). Paraffin tissue blocks were cut into 2-10 µm thick sections using a standard sliding manual microtome (Microm, Dreieich, Germany) and transferred into 1.5 ml sample tubes, washed with xylene (2x speed for 5 min) and absolute ethanol (2x speed for 5 min) and dried. The samples were then homogenised on ice in lysis buffer containing 100 mM Tris-HCl (pH 8.0) and 100 mM dithiothreitol (DTT) and sonicated using Branson sonifier B-12 sonicator for 10-30 s. SDS (4% (w/v), final concentration) was added to the samples and incubated at 99°C for 1 h under mild agitation, cooled to ambient temperature, centrifuged at 2,000 x g for 20 min, and the resulting supernatant fractions were collected [14].

<u>Gastric cancer (GC):</u> The gastric cancer patients underwent a surgical procedure with a complete resection of the tumour. From each donor, tissue samples of tumour mucosa and adjacent normal mucosa were collected and snap-frozen in liquid nitrogen (~50 mg wet weight). The tissue samples were mixed on ice in solubilisation

buffer containing 7 M urea, 2 M thiourea, 40 mM Tris-HCl and a protease inhibitor cocktail mixture (1:100) and homogenised using a Precellys 24 (two cycles at power 5500, 30 s/cycle) in 2 ml reinforced homogenisation tubes (Bertin Technologies, Sweden). The samples were mixed with another round of solubilisation buffer with the addition of 0.5% (w/v) SDS and 10 mM DTT and left for overnight incubation on a shaker at 4°C. After overnight incubation, 25 mM iodoacetamide (IAA) was added and incubated for 40 min in the dark at room temperature. The tissue extracts were centrifuged at 19,500 x g for 20 min. The protein-containing supernatant fractions were diluted ten times with 20 mM sodium bicarbonate, applied to a preconditioned 10 kDa cut-off filter (Pall, Port Washington, USA) and washed three times in the same buffer.

<u>Colorectal cancer (CRC):</u> CRC tumour samples obtained from Sydney South West Area Health Service, Australia were collected from patients undergoing surgical resection [15]. Pathological examination followed the Australian Clinico-Pathological Staging protocol for CRC, which is compatible with other staging systems e.g. the TNM (tumours/nodes/metastases) [3]. All samples were classified as T3N0M0 stage in the TNM classification. The collected CRC tissue samples were stored in Hanks' balanced salt solution at 4°C for approximately 6 h to maintain cell viability. The samples were cut into 2 mm strips and incubated in RPMI 1640 medium containing 2% (v/v) collagenase type 4 (Worthington, USA) and 0.2% (v/v) deoxyribonuclease I from bovine pancreas for 60 min at 37°C on a shaker. The semi-digested tissue was then forced gently through a fine wire mesh strainer using the plunger of a 20 ml syringe and then washed in HBSS. The resulting cell suspension was concentrated in two rounds using 200 µm and 50 µm filters to remove cell aggregates.

The cell suspensions were stored at -80°C in heat-inactivated 90% (v/v) FBS and 10% (v/v) DMSO. Thawed cell suspensions were treated with 0.1% (w/v) deoxyribonuclease I and enriched for epithelial cell fractions by using immuno-magnetic beads conjugated to an antibody against the epithelial molecular marker EpCAM (Dynabead CELLection Epithelial Enrich, Invitrogen). The beads and cells were mixed and incubated for 1 h at 4°C and washed thoroughly with PBS. The proteins were extracted by acetone precipitation (see below).
CRC primary tissue samples obtained from the Department of Pathology in Yonsei University, South Korea, were diagnosed and staged for CRC by a trained pathologist at the Severance Hospital, South Korea. All CRC primary tumours were adenocarcinomas obtained from five male patients spanning differences in age and CRC pathology i.e. varying sites (sigmoid, transverse and rectum), TNM stages (I-IV), and epidermal growth factor receptor (EGFR) expression status [4, 5, 16] (see **Supplementary Table S1 (B)**. The microsomal protein fractions were extracted from the tissue samples as described below.

Prostate cancer (PCa): All biopsy cores were examined for signs of PCa by a trained pathologist. Fresh tissues were collected after radical prostatectomy from a cohort of 55 patients, which included ten patients from each disease stage (PCa grade 1-5) and five individuals presenting with benign prostatic hyperplasia (BPH). The PCa grade was defined using the Modified Gleason Grading System proposed by the International Society of Urological Pathology (ISUP) in 2005, a system that was revised in 2014 [17, 18]. Fresh prostate tissues (~60 mg), stored in RNALater (stabilisation and storage solution), were washed in 80% (v/v) cold ACN and re-suspended in an extraction buffer containing 6 M urea, 10 mM DTT, 1 mM sodium fluoride, 1 mM sodium orthovanadate and a protease inhibitor cocktail (1:10).

Prostate tissues were rapidly lysed (2 min, 30 Hz with a 5 mm stainless steel bead) using a TissueLyser (Qiagen, Chadstone, VIC, Australia). Protein concentrations were determined by the Qubit fluorimetric detection method and directly used for *N*-glycomics.

**Protein extracts from cellular fractions**

As briefly described below, protein extracts from different cellular fractions were obtained from the investigated cell lines and tissue samples.

Secretome (S): Cells grown to high (>95%) confluency were washed at least four times with ice-cold PBS to remove FBS and incubated in FBS-free media at 37°C in 5% $CO_2$ for 28-48 h. The media containing the secreted proteins were collected and centrifuged at 2,000 x g to pellet any cells or cellular debris. The proteins in the supernatant fractions (hereafter the "secretome") were transferred to new vials and concentrated using a 10 kDa molecular weight cut-off Amicon Ultra Centrifugal filter device (Millipore, USA) and stored in 4°C until use.

Whole cell lysates (WCL): Cell pellets (see above) were gently thawed on ice, re-suspended in RIPA lysis buffer, vigorously vortexed and centrifuged at 14,000 x g, 4°C for 10-20 min. The resulting supernatant fractions (the "whole cell lysates") containing the cellular proteins were collected and stored at 4°C until use.

Microsomal fractions (MF): This preparation follows a previously published protocol [19]. In short, cell pellets were re-suspended in 25 mM Tris-HCl (pH 7.4), 150 mM NaCl, and 1 mM EDTA with 5 mM EDTA-free protease inhibitor tablets. The cell suspension was ultra-sonicated using Sonifier 450 on ice over three cycles of 10 s bursts and then centrifuged at 2,000 x g, 4°C for 20 min. The resulting supernatant was ultra-centrifuged at 120,000 x g, 4°C for 80 min. The resulting pellet containing the total membrane complement was re-suspended in 25 mM Tris-HCl (pH 7.4), 150 mM NaCl and 1% (v/v) Triton X-114 and phase partitioned on ice for 10 min, and then at 37°C for 20 min. A subsequent gentle centrifugation step at 1,000 x g at 25°C for 10 min created two distinct phases. The dense detergent phase containing the membrane protein complement (the "microsome") was stored in 4°C until use.

**Protein isolation**

<u>Acetone precipitation:</u> The proteins of all three cellular fractions (WCL, MF, and S, see above) from most cells and tissues investigated in this study were precipitated by adding ice-cold acetone in a ratio of 1:4-1:9 (v/v), vortexed thoroughly and incubated overnight at -20°C in an upright position. The samples were then centrifuged at 14,000 x g for 12-15 min and the protein pellets were stored in -30°C until further analysis.

<u>Chloroform-water-methanol extraction:</u> The proteins in the supernatant fractions after cell lysis of CLL tissues (B cells) and non-melanoma BCC tissues were precipitated according to the method used by Wessel and Flugge [20]. Briefly, the supernatant was mixed vigorously with methanol, chloroform and water in a 4:1:3 volume ratio. The samples were centrifuged at 14,000 x g for 5 min and the upper aqueous phase was carefully removed. Additional methanol was added, mixed and centrifuged as above. The supernatant was removed, and dried protein pellet was stored at -20°C until analysis.

The protein pellets were re-suspended in 6-8 M urea and the protein concentrations were determined using the Bradford assay or the bicinchoninic acid assay.

**Protein denaturation**

Proteins were reduced using 10 mM DTT, 45 min, 56°C, and carbamidomethylated using 25 mM IAA, 30 min in the dark, 20°C before the alkylation reactions were quenched using 30 mM DTT (final concentrations stated).

### Supporting tables

**Supporting Table S1.** Overview of the cultured cell lines, their culture conditions and protein fraction(s) analysed in this study. #Initials of the primary contributor(s) responsible for acquiring the PGC-LC-MS/MS data, see Chatterjee et al., 2019 for details [21]. *Indicates non-cancer cell line; APL: acute promyelocytic leukaemia, AML: acute monocytic leukaemia, ALL: acute lymphocytic leukaemia, S: secretome, WCL: whole cell lysate, MF: microsomal fraction, FF: fresh frozen, FFPE: formalin-fixed and paraffin-embedded, P: paired tissue sample (tumour/non-tumour reference).

| **Cancer type** | **Primary contri-butor#** | **Cell lines (origin/ identifier)** | **Culture media and properties** | **Protein fraction** | **Ref** |
|---|---|---|---|---|---|
| **1. Brain cancer** | | | | | |
| A. Glio-blastoma<br><br>B. Neuro-blastoma | CA | i. U87MG (ATCC HTB-14) | DMEM, adherent | WCL | In prep |
| | IL | i. U87MG[a)] (ATCC HTB-14)<br><br>ii. A172[a)] (ATCC CRL-1620)<br><br>iii. U118MG (CLS 300362)<br><br>iv. U138MG[a)] (CLS 300363) | DMEM, adherent | MF<br><br>MF, S<br>WCL<br><br>MF, WCL<br><br>MF, WCL | In prep |
| | ZSB | i. SK-N-B(2)[b)] (ATCC CRL-2271) | DMEM, adherent | MF | In prep |
| **2. Blood cancer** | | | | | |

| | | | | | |
|---|---|---|---|---|---|
| A. APL | IL | i. HL-60 (ATCC CCL-240) | RPMI 1640, suspension | Differentia-ted/undif-ferrentiated WCL | - |
| B. AML | LYL | i. THP1 (ATCC TIB-202) | RPMI 1640, suspension | Uninfected /infected MF, WCL | [22] |
| C. ALL | MN | i. CCRF-CEM (T-cell) (ATCC CCL-119) | RPMI 1640, suspension | MF | [6] |
| **3. Melanoma skin cancer** | JLA | i. MM253 (CBA-1347) | RPMI 1640, adherent | MF | [8] |
| **4. BC** | LYL | i. HMEC* (ATCC CRL-3243)<br><br>ii. HS578T (ATCC HTB-126)<br><br>iii. MCF7 (ATCC HTB-22)<br><br>iv. MDA-MB157 (ATCC HTB-24)<br><br>v. MDA-MB231 (ATCC HTB-26)<br><br>vi. MDA-MB468 (ATCC HTB-132)<br><br>vii. SKBR3 (ATCC HTB-30) | RPMI 1640, adherent | MF, S | [2] |

| | | | | | |
|---|---|---|---|---|---|
| **5. Lung cancer** | EM | i. A549 (ATCC CCL-185) | RPMI 1640, adherent | MF, WCL | - |
| **6. HCC** | SC | i. HepG2 (ATCC HB-8065) | DMEM, adherent | MF, WCL | - |
| **7. CRC** | JHLC | i. SW480 (ATCC CCL-228)<br><br>ii. SW620 (ATCC CCL-227)<br><br>iii. SW837 (ATCC CCL-235)<br><br>iv. SW1116 (ATCC CCL-233) | DMEM, adherent | MF | [3] |
| | MKS | v. LIM1215[c)]<br><br>vi. LIM1899[c)]<br><br>vii. LIM2405[c)] | RPMI 1640, adherent | MF | [4] |
| **8. OvC** | MA | i. A2780 (ATCC CRL-2772)<br><br>ii. IGROV1 (ATCC TCP-1021)<br><br>iii. OVCAR3 (ATCC HTB-161)<br><br>iv. SKOV3 (ATCC HTB-77)<br>v. HOSE 6.3*[d)] | i-iv) RPMI 1640, adherent<br><br>v-vi) MCDB 105: medium 199, adherent | MF | [9] |

| | | | | | |
|---|---|---|---|---|---|
| | | vi. HOSE 17.1*[d)] | | | |
| **9. BlaCa** | EM | i. J82<br>(ATCC HTB-1)<br><br>ii. RT112<br>(ATCC TCP-1020)<br><br>iii. UROtsa*[e)] | a) MEM alpha media<br><br>b-c) RPMI 1640 adherent | MF, WCL | - |

[a)] Gift from Prof Torsten Pietsch, Institute of Neuropathology, University of Bonn Medical Center, Bonn, Germany

[b)] Gift from Prof Maria Kavallaris, Children's Cancer Institute, Lowy Cancer Research Centre, University of New South Wales, Sydney, NSW, Australia

[c)] Ludwig Institute for Cancer Research, Melbourne, VIC, Australia

[d)] Garvan Institute of Medical Research, Sydney, NSW, Australia

[e)] Gift from Prof Philip Erben, Department of Urology, Medical Faculty Mannheim, Heidelberg University, Mannheim, Germany

**Supporting Table S2.** Details of cultured cells investigated in this study. *Indicates non-cancer cell lines; M: male, F: female, APL: acute promyelocytic leukaemia, AML: acute monocytic leukaemia, ALL: acute lymphocytic leukaemia.

| Cancer type | Cell lines (origin/ identifier) | Age (years) | Sex | Source | Tumourigenic |
|---|---|---|---|---|---|
| **1. Brain cancer** | | | | | |
| A. Glio-blastoma | i. U87MG[a)] (ATCC HTB-14) | - | M | Glioblastoma | Yes |
| | ii. A172 (ATCC CRL-1620) | 53 | M | Glioblastoma | No |
| | iii. U118MG[a)] (CLS 300362) | 50 | M | Glioblastoma | Yes |
| | iv. U138MG[a)] (CLS 300363) | 47 | M | Glioblastoma | Yes |
| B. Neuro-blastoma | i. SK-N-B(2)[b)] (ATCC CRL-2271) | 2 | M | Neuroblastoma | Yes |
| **2. Blood cancer** | | | | | |
| A. APL | i. HL-60 (ATCC CCL-240) | 36 | F | Promyelocytes from peripheral blood | Yes |
| B. AML | i. THP1 (ATCC TIB-202) | 1 | M | Monocytes from peripheral blood | No |
| C. ALL | i. CCRF-CEM (ATCC CCL-119) | 4 | F | T-lymphoblasts from peripheral blood | Yes |
| **3. Melanoma skin cancer** | i. MM253 (CBA-1347) | - | - | Lymph node | Yes |
| **4. BC** | i. HMEC* | | | | |

| | | | | | |
|---|---|---|---|---|---|
| | (ATCC PCS600-010 | - | F | Cuboidal epithelium BC | No |
| | ii. HS578T (ATCC HTB-126) | 74 | F | Sarcoma (Basal B) | No |
| | iii. MCF7 (ATCC HTB-22) | 69 | F | Pleural effusion, luminal adeno-carcinoma | No |
| | iv. MDA-MB157 (ATCC HTB-24) | 44 | F | Pleural effusion, luminal adeno-carcinoma | Yes |
| | v. MDA-MB231 (ATCC HTB-26) | 51 | F | Pleural effusion, metastatic adenocarcinoma (Basal B) | Yes |
| | vi. MDA-MB468 (ATCC HTB-132) | 51 | F | Pleural effusion, Luminal adeno-carcinoma | Yes |
| | vii. SKBR3 (ATCC HTB-30) | 43 | F | Pleural effusion, HER2-enriched adeno-carcinoma | Yes |
| **5. Lung cancer** | i. A549 (ATCC CCL-185) | 58 | M | Lung tissues | Yes |
| **6. HCC** | i. HepG2 (ATCC HB-8065) | 15 | M | HCC tissues | Yes |
| **7. CRC** | i. SW480 (ATCC CCL-228) | 50 | M | Dukes' Type B CRC (rectum) | Yes |
| | ii. SW620 (ATCC CCL-227) | 51 | M | Dukes' Type C CRC (Colon) | Yes |
| | iii. SW837 (ATCC CCL-235) | 53 | M | Dukes' Type C CRC (Rectum) | Yes |
| | iv. SW1116 | 73 | M | Dukes' Type A | |

| | | | | | |
|---|---|---|---|---|---|
| | (ATCC CCL-233) | | | CRC (Large Intestine, Colon) | Yes |
| | v. LIM1215[c)] | 34 | M | Hereditary nonpolyposis CRC, Ileocecal valve, Omental metastasis | Yes |
| | vi. LIM1899 [c)] | - | - | Columnar cell carcinoma of invasive colon, moderately differentiated | Yes |
| | vii. LIM2405 [c)] | - | M | Columnar cell carcinoma of invasive colon, poorly differentiated | Yes |
| **8. OvC** | i. A2780 (ATCC CRL-2772) | - | F | Serous (ovary) | Yes |
| | ii. IGROV1 (ATCC TCP-1021) | 47 | F | Endometroid and serous (right ovary) | Yes |
| | iii. OVCAR3 (ATCC HTB-161) | 60 | F | Serous (ovary, ascites fluid) | Yes |
| | iv. SKOV3 (ATCC HTB-77) | 64 | F | Serous (ovary, ascites fluid) | Yes |
| | v. HOSE 6.3*[d)] | - | F | Normal ovary | No |
| | vi. HOSE 17.1*[d)] | - | F | Normal ovary (HOSE 6.3 and 17.1 are infected with Human papillomavirus type 16) | No |

| | | | | | |
|---|---|---|---|---|---|
| **9. BlaCa** | i. J82 (ATCC HTB-1) | 58 | M | Transitional cell carcinoma (basal) | Yes |
| | ii. RT112 (ATCC TCP-1020) | - | F | Transitional cell carcinoma (luminal) | Yes |
| | iii. UROtsa*[e)] | 12 | F | “Normal” urothelium immortalised using SV40 large T-antigen | No |

[a)] Gift from Professor Torsten Pietsch, Institute of Neuropathology, University of Bonn Medical Center, Bonn, Germany

[b)] Gift from Prof Maria Kavallaris, Children’s Cancer Institute, Lowy Cancer Research Centre, University of New South Wales, Sydney, NSW, Australia

[c)] Ludwig Institute for Cancer Research, Parkville, VIC, Australia

[d)] Garvan Institute of Medical Research, Sydney, NSW, Australia

[e)] Gift from Professor Philip Erben, Department of Urology, Medical Faculty Mannheim, Heidelberg University, Mannheim, Germany

**Supporting Table S3.** Overview of the tissue samples investigated in this study and relevant ethics details. #Initials of the primary contributor responsible for acquiring the PGC-LC-MS/MS data, see Chatterjee et al., 2019 for details [21]. *Indicates non-cancer cell lines; S: secretome, WCL: whole cell lysate, MF: microsomal fraction, FF: fresh frozen, FFPE: formalin-fixed and paraffin-embedded, P: paired tissue samples (tumour vs non-tumour reference).

| Cancer type | Primary contri-butor# | Biological replicates (n) | Sample type | Ethics details | Ref |
|---|---|---|---|---|---|
| **1. Blood cancer** | | | | | |
| A. CLL | KS | i. 8 | FF | Project #8935 University of Sydney, Sydney, Australia | - |
| **2. Non-melanoma skin cancer** | | | | | |
| A. BCC<br>B. SCC | UM | i. 14<br>ii. 20<br>i. 15 | FF BCC (P)<br>FFPE BCC (P)<br>FFPE SCC (P) | 127-11-18042011 University of Leipzig, Leipzig, Germany | [1] |
| **3. GC** | VV | i. 3 | FF (P) | Comitato Etico Regione Toscana (Tuscany, Italy) | - |
| **4. HCC** | HH | i. 3 | FFPE (P) | Klinieka Bolnica Merkur (Zagreb, Croatia) | [7] |
| **5. CRC** | JHLC | i. 6 | FF (P) | X08-0614 Sydney South West | [3] |

| | | | | | |
|---|---|---|---|---|---|
| | | | | Area Health Service, Australia | |
| | MKS | ii. 5 | FF (P) | 5201100040 Yonsei University, Seoul, South Korea | [4] |
| **6. PCa** | RKS | i. 50 tumour 5 BPH | FF | nº2695126 Faculdade de Medicina do Estado de São Paulo, Brazil | In prep |

**Supporting Table S4:** Details of individual tissue samples investigated in this study. M: male, F: female, FF: fresh frozen, FFPE: formalin-fixed and paraffin-embedded, P: paired tissue sample (tumour/non-tumour reference).

| **Cancer type** | **Patient #** | **Age** (years) | **Sex** | **Loca-lisation** | **Tumour invasion level$** | **Diff. status / doubling time** |
|---|---|---|---|---|---|---|
| **1. Blood Cancer** | | | | | | |
| A. CLL | FF 1 | - | - | Blood | Stable | > 5 years |
| | FF 2 | - | - | Blood | Inter-mediate | < 18 months |
| | FF 3 | - | - | Blood | Inter-mediate | < 18 months |
| | FF 4 | - | - | Blood | Inter-mediate | < 18 months |
| | FF 5 | - | - | Blood | Inter-mediate | < 18 months |
| | FF 6 | - | - | Blood | Progressive | < 6 months |
| | FF 7 | - | - | Blood | Progressive | < 6 months |
| | FF 8 | - | - | Blood | Progressive | < 6 months |
| **2. Non-melanoma skin cancer** | | | | | | |
| **A. BCC** | FF 1 | 74 | M | Lower Leg | IV | - |
| | FF 2 | 79 | M | Shoulder | IV | - |
| | FF 3 | 79 | M | Shoulder | IV | - |
| | FF 4 | 74 | M | Lower leg | IV | - |
| | FF 5 | 73 | M | Back | IV | - |
| | FF 6 | 72 | M | Chest | IV | - |
| | FF 7 | 82 | M | Shoulder | IV | - |
| | FF 8 | 90 | M | Back | III | - |
| | FF 9 | 88 | M | Shoulder | IV | - |
| | FF 10 | 80 | F | Cheek | IV | - |
| | FF 11 | 70 | M | Chest | IV | - |

| | | | | | | |
|---|---|---|---|---|---|---|
| | FF 12 | 82 | M | Shoulder | IV | - |
| | FF 13 | 75 | M | Shoulder | IV | - |
| | FF 14 | 80 | M | Shoulder | IV | - |
| | FFPE 1 | 71 | M | Nose | IV | - |
| | FFPE 2 | 25 | F | Ear | III | - |
| | FFPE 3 | 88 | M | Nose | IV | - |
| | FFPE 4 | 25 | F | Head | III | - |
| | FFPE 5 | 79 | M | Back | III | - |
| | FFPE 6 | 79 | M | Neck | IV | - |
| | FFPE 7 | 72 | F | Cheek | IV | - |
| | FFPE 8 | 75 | F | Nose | IV | - |
| | FFPE 9 | 90 | M | Chest | IV | - |
| | FFPE 10 | 62 | M | Nose | IV | - |
| | FFPE 11 | 73 | F | Nose | IV | - |
| | FFPE 12 | 82 | F | Nose | IV | - |
| | FFPE 13 | 72 | M | Cheek | IV | - |
| | FFPE 14 | 62 | M | Shoulder | IV | - |
| | FFPE 15 | 77 | F | Forehead | IV | - |
| | FFPE 16 | 80 | F | Chin | IV | - |
| | FFPE 17 | 84 | M | Lower leg | IV | - |
| | FFPE 18 | 81 | M | Cheek | IV | - |
| | FFPE 19 | 77 | M | Cheek | IV | - |
| | FFPE 20 | 58 | M | Nose | V | - |
| **B. SCC** | FFPE 1 | 65 | F | Cheek | IV | - |
| | FFPE 2 | 85 | M | Head | V | - |
| | FFPE 3 | 72 | F | Head | IV | - |
| | FFPE 4 | 90 | M | Head | V | - |
| | FFPE 5 | 72 | M | Nose | V | - |
| | FFPE 6 | 74 | F | Cheek | IV | - |
| | FFPE 7 | 73 | F | Upper trunk | III | - |
| | FFPE 8 | 76 | M | Cheek | IV | - |
| | FFPE 9 | 74 | M | Head | IV | - |
| | FFPE 10 | 79 | F | Head | IV | - |

| | | | | | | |
|---|---|---|---|---|---|---|
| | FFPE 11 | 90 | M | Head | V | - |
| | FFPE 12 | 88 | M | Ear | IV | - |
| | FFPE 13 | 64 | M | Head | IV | - |
| | FFPE 14 | 87 | M | Ear | V | - |
| | FFPE 15 | 78 | M | Head | V | - |
| **3. GC** | FF 1 | 70 | M | Gastric | G3 | |
| | FF 2 | 64 | M | Gastric | NG | |
| | FF 3 | 76 | F | Gastric | G2 | |
| **4. HCC** | - | - | - | - | - | - |
| **5. CRC** | | | | | | |
| **JHLC** | FF 1 | 76 | F | Rectum | B1/T3N0M0 | Moderate |
| | FF 2 | 84 | F | Colon | B1/T3N0M0 | Moderate |
| | FF 3 | 56 | F | Rectum | B1/T3N0M0 | Moderate |
| | FF 4 | 57 | M | - | B1/T3N0M0 | Moderate |
| | FF 5 | 74 | F | - | B1/T3N0M0 | Moderate |
| | FF 6 | - | - | - | - | - |
| **MKS** | FF 1 | 64 | M | Sigmoid, polyploid | I | Well |
| | FF 2 | 69 | M | Rectum, ulcerative | IIIB | Moderate |
| | FF 3 | 67 | M | Cecum, ulcero fungating | IIIB | Moderate |
| | FF 4 | 40 | M | Rectum, annular constrictive | IV | Well |
| | FF 5 | 73 | M | Transverse, ulcero fungating | IV | Poor |
| **6. PCa** | FF 1 | 62 | M | Prostate | 3/3/1 (T1c) | - |
| | FF 2 | 68 | M | Prostate | 3/3/1 (T3a) | - |
| | FF 3 | 54 | M | Prostate | 3/3/1 (T2b) | - |

| | | | | | | |
|---|---|---|---|---|---|---|
| | FF 4 | 60 | M | Prostate | 3/3/1 (T1c) | - |
| | FF 5 | 52 | M | Prostate | 3/3/1 (T1c) | - |
| | FF 6 | 52 | M | Prostate | 3/3/1 (T1c) | - |
| | FF 7 | 59 | M | Prostate | 3/3/1 (T1c) | - |
| | FF 8 | 59 | M | Prostate | 3/3/1 (T2c) | - |
| | FF 9 | 64 | M | Prostate | 3/3/1 (T2a) | - |
| | FF 10 | 67 | M | Prostate | 3/3/1 (T1c) | - |
| | FF 11 | 67 | M | Prostate | 3/4/2 (T2b) | - |
| | FF 12 | 72 | M | Prostate | 3/4/2 (T1c) | - |
| | FF 13 | 50 | M | Prostate | 3/4/2 (T1c) | - |
| | FF 14 | 72 | M | Prostate | 3/4/2 (T2b) | - |
| | FF 15 | 63 | M | Prostate | 3/4/2 (T1c) | - |
| | FF 16 | 68 | M | Prostate | 3/4/2 (T1c) | - |
| | FF 17 | 58 | M | Prostate | 3/4/2 (T1c) | - |
| | FF 18 | 68 | M | Prostate | 3/4/2 (T1c) | - |
| | FF 19 | 66 | M | Prostate | 3/4/2 (T1c) | - |
| | FF 20 | 57 | M | Prostate | 3/4/2 (T1c) | - |

| | | | | | | |
|---|---|---|---|---|---|---|
| | FF 21 | 79 | M | Prostate | 4/3/3 (T1c) | - |
| | FF 22 | 55 | M | Prostate | 4/3/3 (T1c) | - |
| | FF 23 | 70 | M | Prostate | 4/3/3 (T2c) | - |
| | FF 24 | 64 | M | Prostate | 4/3/3 (T1c) | - |
| | FF 25 | 66 | M | Prostate | 4/3/3 (T1c) | - |
| | FF 26 | 62 | M | Prostate | 4/3/3 (T2b) | - |
| | FF 27 | 70 | M | Prostate | 4/3/3 (T1c) | - |
| | FF 28 | 62 | M | Prostate | 4/3/3 (T1c) | - |
| | FF 29 | 55 | M | Prostate | 4/3/3 (T2a) | - |
| | FF 30 | 48 | M | Prostate | 4/3/3 (T2a) | - |
| | FF 31 | 71 | M | Prostate | 4/4/4 (T2b) | - |
| | FF 32 | 69 | M | Prostate | 4/4/4 (T2b) | - |
| | FF 33 | 56 | M | Prostate | 4/4/4 (T2c) | - |
| | FF 34 | 50 | M | Prostate | 4/4/4 (T2c) | - |
| | FF 35 | 66 | M | Prostate | 4/4/4 (T2c) | - |
| | FF 36 | 75 | M | Prostate | 4/4/4 (T2b) | - |
| | FF 37 | 60 | M | Prostate | 4/4/4 (T2b) | - |

| | | | | | | |
|---|---|---|---|---|---|---|
| | FF 38 | 66 | M | Prostate | 4/4/4 (T1c) | - |
| | FF 39 | 63 | M | Prostate | 4/4/4 (T1c) | - |
| | FF 40 | 63 | M | Prostate | 4/4/4 (T1c) | - |
| | FF 41 | 60 | M | Prostate | 5/5/5 (T2c) | - |
| | FF 42 | 68 | M | Prostate | 5/5/5 (T2b) | - |
| | FF 43 | 56 | M | Prostate | 5/5/5 (T2c) | - |
| | FF 44 | 59 | M | Prostate | 5/5/5 (T2c) | - |
| | FF 45 | 66 | M | Prostate | 5/5/5 (T2c) | - |
| | FF 46 | 65 | M | Prostate | 5/5/5 (T2a) | - |
| | FF 47 | 70 | M | Prostate | 5/5/5 (T3c) | - |
| | FF 48 | 73 | M | Prostate | 5/5/5 (T2c) | - |
| | FF 49 | 58 | M | Prostate | 5/5/5 (T1c) | - |
| | FF 50 | 65 | M | Prostate | 5/5/5 (T1c) | - |
| | FF 51 | 3.5 | M | Prostate | BPH | - |
| | FF 52 | 10.2 | M | Prostate | BPH | - |
| | FF 53 | 6.9 | M | Prostate | BPH | - |
| | FF 54 | 4.4 | M | Prostate | BPH | - |
| | FF 55 | 2.3 | M | Prostate | BPH | - |

$Please note that the annotations of the tumour grade and invasion level, which were performed by trained pathologists across multiple institutions, vary between the investigated cancer types and subtypes.

**Supporting Table S5.** Details of formalin-fixed and paraffin-embedded (FFPE) tissue microarray (TMA) samples from patients diagnosed with cancers investigated by MALDI-MSI in this study. M: male, F: female.

| **Organ** | **Diagnosis** | **Age** (years) | **Sex** | **Tumour history** | **Differentiation / Tumour grade** |
|---|---|---|---|---|---|
| **Skin** | Cancer (Bi-ii, C) | 43 | M | N/A | N/A |
| | Normal | N/A | F | - | - |
| **Colon-rectum** | CRC (Di-ii) | 54 | M | 7 months | Moderate/ T3N0M0 |
| | Normal | 24 | M | - | - |
| **Liver** | HCC (F) | 65 | M | 15 days | Moderate/ T1N0M0 |
| | Normal | 30 | M | - | - |
| **Gastric / Stomach** | GC (G) | 48 | M | 1 month | Poor / T3N1M0 |
| | Normal | 66 | M | - | - |

**Supporting Table S6.** Distribution (relative abundance) of the *N*-glycan types across various protein fractions (WCL, MF, S) extracted from the studied human cancer cell lines.

| Cancer type | Cell Lines / sample type | Distribution of *N*-glycan types | | |
|---|---|---|---|---|
| | | **Pauci-mannose** | **Oligo-mannose** | **Complex/ hybrid** |
| **1. Brain cancer** | | | | |
| A. Glioblastoma | i. U87MG_MF<br>U87MG _WCL | 6.87%<br>14.34% | 35.19%<br>47.70% | 57.94%<br>37.96% |
| | ii. A172_MF<br>A172_S<br>A172_WCL | 13.25%<br>1.20%<br>29.97% | 58.36%<br>8.19%<br>49.69% | 28.39%<br>90.61%<br>20.34% |
| | iii. U118MG_MF<br>U118MG_WCL | 6.82%<br>4.92% | 38.08%<br>20.73% | 55.10%<br>74.35% |
| | iv. U138MG_MF<br>U138MG _WCL | 7.51%<br>8.21% | 61.73%<br>26.12% | 30.76%<br>65.67% |
| B. Neuro-blastoma | i. SK-N-B(2) | 1.80% | 59.91% | 38.29% |
| **2. Blood cancer** | | | | |
| A. APL | i. HL60_WCL<br>Differentiated<br>Undifferentiated | <br>19.91%<br>2.68% | <br>77.15%<br>91.21% | <br>2.58%<br>6.11% |
| B. AML | i. THP1_MF<br>Infected<br>Uninfected<br>THP1_WCL<br>Infected<br>Uninfected | <br>11.76%<br>11.82%<br><br>40.22%<br>50.20% | <br>37.61%<br>39.19%<br><br>36.21%<br>37.71% | <br>50.63%<br>48.99%<br><br>23.57%<br>12.09% |
| C. ALL | i. CCRF-CEM_MF | 0.00% | 34.30% | 65.70% |
| **3. Melanoma skin cancer** | i. MM253 | 6.74% | 68.60% | 24.66% |
| | i. HMEC*_MF<br>HMEC*_S | 0.00%<br>0.00% | 64.38%<br>6.11% | 35.62%<br>93.89% |

| | | | | |
|---|---|---|---|---|
| **4. BC** | ii. HS578T_MF | 0.00% | 51.39% | 48.61% |
| | HS578T_S | 0.00% | 5.28% | 94.72% |
| | iii. MCF7_MF | 3.28% | 80.88% | 15.84% |
| | MCF7_S | 0.89% | 23.82% | 75.29% |
| | iv. MDA-B157_MF | 11.59% | 50.28% | 38.13% |
| | MDA-MB157_S | 3.21% | 12.90% | 83.89% |
| | v. MDA_MB231_MF | 0.29% | 44.66% | 55.05% |
| | MDA-MB231_S | 9.49% | 11.41% | 79.10% |
| | vi. MDA_MB468_MF | 11.33% | 80.26% | 8.41% |
| | MDA-MB468_S | 2.07% | 10.39% | 87.54% |
| | vii. SKBR3_MF | 6.18% | 52.55% | 41.27% |
| | SKBR3_S | 1.03% | 15.48% | 83.49% |
| **5. Lung cancer** | i. A549_MF | 5.68% | 71.29% | 23.03% |
| | A549_WCL | 5.01% | 72.27% | 22.72% |
| **6. HCC** | i. HepG2_MF | 6.69% | 59.10% | 34.21% |
| | HepG2_WCL | 18.85% | 67.35% | 13.79% |
| **7. CRC** | i. SW480_MF | 1.38% | 56.45% | 42.17% |
| | ii. SW620_MF | 2.77% | 50.45% | 46.78% |
| | iii. SW837_MF | 1.55% | 60.41% | 38.05% |
| | iv. SW1116_MF | 1.52% | 50.51% | 47.97% |
| | v. LIM1215_MF | 16.74% | 72.88% | 10.37% |
| | vi. LIM1899_MF | 19.62% | 68.70% | 11.68% |
| | vii. LIM2405_MF | 20.31% | 58.93% | 20.75% |
| **8. OvC** | i. A2780_MF | 10.20% | 57.34% | 32.47% |
| | ii. IGROV1_MF | 3.69% | 61.78% | 34.53% |
| | iii. OVCAR3_MF | 3.56% | 63.09% | 33.35% |
| | iv. SKOV3_MF | 1.96% | 58.72% | 39.32% |
| | v. HOSE 6.3_MF | 1.49% | 21.28% | 77.22% |
| | vi. HOSE 17.1_MF | 2.64% | 37.93% | 59.43% |
| **9. BlaCa** | i. J82_MF | 2.86% | 41.53% | 55.62% |
| | J82_WCL | 2.50% | 41.27% | 56.23% |
| | ii. RT112_MF | 1.95% | 46.46% | 51.60% |
| | RT112_WCL | 1.98% | 57.45% | 40.57% |
| | iii. UROtsa*_MF | 3.02% | 62.88% | 34.09% |
| | UROtsa*_ WCL | 2.33% | 49.24% | 48.43% |

**Supporting Table S7.** Distribution (relative abundance) of the individual oligomannosidic species across various protein fractions (WCL, MF, S) extracted from the studied human cancer cell lines.

| Cancer type | Cell lines_ Sample type | Distribution of Oligomannose | | | | |
|---|---|---|---|---|---|---|
| | | M5 | M6 | M7 | M8 | M9 |
| **1. Brain cancer** | | | | | | |
| A. Glioblastoma | i. U87MG_MF | 2.26% | 5.25% | 5.92% | 9.57% | 12.19% |
| | U87MG _WCL | 6.32% | 11.22% | 10.14% | 11.15% | 8.87% |
| | ii. A172_MF | 1.87% | 6.47% | 16.01% | 18.62% | 15.39% |
| | A172_S | 2.45% | 0.98% | 1.66% | 2.01% | 1.09% |
| | A172_WCL | 3.42% | 6.37% | 10.61% | 13.80% | 15.50% |
| | iii. U118MG_MF | 1.20% | 4.74% | 5.70% | 13.17% | 13.27% |
| | U118MG_WCL | 1.26% | 3.02% | 2.64% | 6.15% | 7.66% |
| | iv. U138MG_MF | 2.12% | 7.09% | 9.28% | 20.22% | 23.01% |
| | U138MG _WCL | 1.72% | 3.93% | 3.95% | 7.59% | 8.93% |
| B. Neuroblastoma | i. SK-N-B(2) | 3.02% | 4.57% | 4.92% | 11.52% | 35.88% |
| **2. Blood cancer** | | | | | | |
| A. APL | i. HL60_WCL | | | | | |
| | Differentiated | 5.83% | 13.39% | 23.35% | 22.60% | 22.26% |
| | Undifferentiated | 2.52% | 12.76% | 12.86% | 32.85% | 30.22% |
| B. AML | i. THP1_MF | | | | | |
| | Infected | 5.49% | 4.80% | 6.42% | 8.95% | 13.52% |
| | Uninfected | 4.89% | 3.91% | 8.61% | 6.84% | 15.93% |
| | THP1_WCL | | | | | |
| | Infected | 3.50% | 3.38% | 7.02% | 9.09% | 13.22% |
| | Uninfected | 3.82% | 3.95% | 9.71% | 7.64% | 15.06% |
| C. ALL | i. CCRF-CEM_MF | 0.12% | 0.00% | 5.26% | 11.23% | 17.68% |
| **3. Melanoma skin cancer** | i. MM253 | 3.28% | 11.24% | 18.74% | 16.63% | 18.71% |
| **4. BC** | i. HMEC*_MF | 0.00% | 8.23% | 11.12% | 22.66% | 22.36% |
| | HMEC*_S | 0.61% | 1.97% | 3.52% | 0.00% | 0.00% |
| | ii. HS578T_MF | 0.00% | 6.56% | 4.58% | 18.77% | 21.47% |

| | | | | | | |
|---|---|---|---|---|---|---|
| | HS578T_S | 0.00% | 1.23% | 1.43% | 2.21% | 0.41% |
| | iii. MCF7_MF | 2.98% | 10.29% | 10.65% | 28.40% | 28.56% |
| | MCF7_S | 4.26% | 5.20% | 6.08% | 4.94% | 3.35% |
| | iv. MDA-MB157_MF | 3.30% | 6.90% | 9.66% | 13.73% | 16.69% |
| | MDA-MB157_S | 1.40% | 3.27% | 3.19% | 3.55% | 1.49% |
| | v. MDA_MB231_MF | 0.36% | 2.82% | 4.69% | 7.78% | 29.01% |
| | MDA-MB231_S | 1.01% | 2.10% | 3.45% | 3.43% | 1.43% |
| | vi. MDA_MB468_MF | 7.41% | 9.77% | 7.68% | 22.27% | 33.13% |
| | MDA-MB468_S | 1.85% | 2.18% | 2.70% | 2.04% | 1.61% |
| | vii. SKBR3_MF | 3.00% | 9.42% | 9.03% | 16.13% | 14.97% |
| | SKBR3_S | 0.91% | 3.61% | 4.12% | 4.20% | 2.64% |
| **5. Lung cancer** | i. A549_MF | 1.00% | 19.27% | 17.00% | 20.08% | 13.94% |
| | A549_WCL | 1.02% | 19.16% | 18.05% | 21.09% | 12.95% |
| **6. HCC** | i. HepG2_MF | 5.60% | 7.46% | 6.77% | 18.39% | 20.88% |
| | HepG2_WCL | 12.86% | 8.44% | 9.82% | 19.48% | 16.75% |
| **7. CRC** | i. SW480_MF | 1.46% | 3.07% | 9.29% | 2.11% | 40.52% |
| | ii. SW620_MF | 4.63% | 6.22% | 12.47% | 24.51% | 2.62% |
| | iii. SW837_MF | 0.61% | 7.07% | 14.63% | 17.56% | 20.55% |
| | iv. SW1116_MF | 0.86% | 5.09% | 10.57% | 15.43% | 18.55% |
| | v. LIM1215_MF | 2.93% | 22.99% | 45.67% | 0.25% | 1.04% |
| | vi. LIM1899_MF | 1.17% | 4.19% | 8.18% | 16.01% | 39.16% |
| | vii. LIM2405_MF | 2.98% | 18.98% | 31.57% | 0.55% | 4.86% |
| **8. OvC** | i. A2780_MF | 1.59% | 4.78% | 12.17% | 17.07% | 21.73% |
| | ii. IGROV1_MF | 0.60% | 7.64% | 13.01% | 15.48% | 25.06% |
| | iii. OVCAR3_MF | 2.14% | 4.75% | 11.18% | 18.86% | 26.16% |
| | iv. SKOV3_MF | 0.69% | 4.62% | 8.81% | 18.85% | 25.76% |
| | v. HOSE 6.3_MF | 0.29% | 1.85% | 4.63% | 7.40% | 7.10% |
| | vi. HOSE 17.1_MF | 0.63% | 2.85% | 6.13% | 13.82% | 14.49% |
| **9. BlaCa** | i. J82_MF | 3.89% | 8.65% | 7.27% | 15.06% | 6.65% |
| | J82_WCL | 3.38% | 9.45% | 6.26% | 15.71% | 6.48% |
| | ii. RT112_MF | 2.16% | 12.37% | 8.93% | 15.02% | 7.97% |
| | RT112_WCL | 2.85% | 17.84% | 10.16% | 14.69% | 11.92% |
| | iii. UROtsa_MF | 3.50% | 17.83% | 11.50% | 14.90% | 15.16% |
| | UROtsa_WCL | 2.68% | 12.79% | 9.59% | 15.66% | 8.51% |

**Supporting Table S8.** Distribution (relative abundance) of the *N*-glycan types across various sample types (FF, FFPE) extracted from the studied human cancer tumour vs non-tumour tissues.

| Cancer type | Sample type_ Biological replicates | Distribution of *N*-glycan types | | |
|---|---|---|---|---|
| | | Pauci-mannose | Oligo-mannose | Complex/ hybrid |
| **1. Blood cancer** | | | | |
| A. CLL | i. FF_Tumour Early stage_8 | 2.13% | 65.57% | 32.30% |
| | FF_Tumour Late stage_8 | 3.37% | 61.82% | 34.81% |
| **2. Non-melanoma skin cancer** | | | | |
| A. BCC | i. FF_Non tumour_14 | 0.88% | 5.21% | 93.91% |
| | FF_Tumour_14 | 1.53% | 9.57% | 88.90% |
| | ii. FFPE_Non tumour_20 | 3.25% | 8.85% | 87.90% |
| | FFPE_Tumour_20 | 3.85% | 10.66% | 84.18% |
| B. SCC | i. FFPE_Non tumour_15 | 2.66% | 8.68% | 88.66% |
| | FFPE_Tumour_15 | 3.89% | 13.10% | 83.01% |
| **3. GC** | i. FF_Non tumour_3 | 1.36% | 23.96% | 74.68% |
| | FF_Tumour_3 | 3.35% | 21.62% | 75.03% |
| **4. HCC** | i. FFPE_Non tumour_3 | 4.78% | 30.31% | 64.91% |
| | FFPE_Tumour_3 | 29.00% | 27.51% | 43.49% |
| **5. CRC** | i. FF_Non tumour_6 | 1.24% | 27.27% | 71.49% |
| | FF_Tumour_6 | 2.72% | 39.31% | 57.96% |
| | ii. FF_Non tumour_5 | 3.52% | 29.68% | 66.80% |
| | FF_Tumour_5 | 5.65% | 38.89% | 55.46% |
| **6. PCa** | i. FF_Non tumour_5 | 3.06% | 12.58% | 84.36% |
| | FF_Tumour_Stage1_10 | 5.12% | 12.26% | 82.62% |
| | FF_Tumour_Stage2_10 | 5.48% | 12.27% | 82.26% |
| | FF_Tumour_Stage3_10 | 6.53% | 11.71% | 81.76% |
| | FF_Tumour_Stage4_10 | 6.68% | 11.62% | 81.71% |
| | FF_Tumour_Stage5_10 | 4.50% | 10.81% | 85.32% |

**Supporting Table S9.** Distribution (relative abundance) of the individual oligomannosidic species across various sample types (FF, FFPE) extracted from the studied human cancer tumour vs non-tumour tissues.

| Cancer type | Sample type_ Biological replicates | Distribution of *N*-glycan types | | | | |
|---|---|---|---|---|---|---|
| | | **M5** | **M6** | **M7** | **M8** | **M9** |
| **1. Blood cancer** | | | | | | |
| A. CLL | i. FF_Tumour Early stage_8 | 1.41% | 8.05% | 11.91% | 18.26% | 25.95% |
| | FF_Tumour Late stage_8 | 1.86% | 6.72% | 12.42% | 21.38% | 19.45% |
| **2. Non-melanoma skin cancer** | | | | | | |
| A. BCC | i. FF_Non tumour_14 | 1.41% | 1.27% | 0.79% | 0.92% | 0.82% |
| | FF_Tumour_14 | 1.93% | 2.28% | 1.61% | 2.02% | 1.73% |
| | ii. FFPE_Non tumour_20 | 1.49% | 1.76% | 1.78% | 2.09% | 1.74% |
| | FFPE_Tumour_20 | 1.54% | 2.18% | 1.41% | 0.78% | 4.75% |
| B. SCC | i. FFPE_Non tumour_15 | 1.97% | 1.75% | 1.41% | 1.89% | 1.67% |
| | FFPE_Tumour_15 | 2.10% | 2.44% | 2.14% | 3.49% | 2.93% |
| **3. GC** | i. FF_Non tumour_3 | 4.97% | 2.80% | 1.67% | 5.84% | 8.67% |
| | FF_Tumour_3 | 4.40% | 3.89% | 3.29% | 5.06% | 4.99% |
| **4. HCC** | i. FFPE_Non tumour_3 | 2.62% | 7.56% | 9.73% | 6.73% | 3.67% |
| | FFPE_Tumour_3 | 8.35% | 7.16% | 7.32% | 2.73% | 1.95% |
| **5. CRC** | i. FF_Non tumour_6 | 3.14% | 2.90% | 4.04% | 6.04% | 11.14% |
| | FF_Tumour_6 | 4.13% | 3.08% | 6.54% | 8.93% | 16.63% |
| | ii. FF_Non tumour_5 | 6.30% | 4.31% | 4.29% | 5.67% | 9.12% |
| | FF_Tumour_5 | 8.45% | 7.81% | 5.69% | 7.08% | 9.85% |
| **6. PCa** | i. FF_Non tumour_5 | 6.50% | 3.18% | 1.19% | 1.17% | 0.53% |
| | FF_Tumour_Stage1_10 | 6.29% | 3.14% | 1.30% | 1.05% | 0.47% |
| | FF_Tumour_Stage2_10 | 6.60% | 2.99% | 1.24% | 0.98% | 0.46% |
| | FF_Tumour_Stage3_10 | 6.14% | 3.01% | 1.14% | 0.95% | 0.47% |
| | FF_Tumour_Stage4_10 | 5.78% | 3.00% | 1.21% | 1.08% | 0.55% |
| | FF_Tumour_Stage5_10 | 4.98% | 2.67% | 1.14% | 0.92% | 0.48% |
